# InferAging: A non-invasive aging clock for quantifying individual differences in aging

**DOI:** 10.64898/2026.05.22.727332

**Authors:** Yoshihiro Ando, Yuichiro Yada, Makoto Kashima, Yasumasa Bessho, Hiromi Hirata, Honda Naoki, Takaaki Matsui

**Affiliations:** Laboratory of Biosystem Dynamics, Division of Biological Sciences, Graduate School of Science and Technology, Nara Institute of Science and Technology (NAIST), Ikoma, Nara, Japan; Laboratory for Data-driven Biology, Nagoya University Graduate School of Medicine, Nagoya, Aichi, Japan; Institute for Advanced Research, Nagoya University, Nagoya, Aichi, Japan; Department of Biomolecular Science, Faculty of Science, Toho University, Funabashi, Chiba, Japan; Laboratory of Gene Regulation Research, NAIST, Ikoma, Nara, Japan; Laboratory of Brain Science, Department of Chemistry and Biological Science, College of Science and Engineering, Aoyama Gakuin University, Sagamihara, Kanagawa, Japan; Laboratory of Data-driven Biology, Graduate School of Integrated Sciences for Life, Hiroshima University, Higashi-hiroshima, Hiroshima, Japan; Center for One Medicine Innovative Translational Research (COMIT), Nagoya University, Showa-ku, Nagoya, Aichi, 466-8550, Japan; Life Science Collaboration Center (LiSCo), NAIST, Ikoma, Nara, Japan; Medilux Research Center, NAIST, Ikoma, Nara, Japan; Center for ARWIT Promotion, NAIST, Ikoma, Nara, Japan

## Abstract

Quantifying differences in aging among individuals of the same chronological age could provide a direct measure of biological aging. However, existing methods often estimate biological age by predicting chronological age and treat such individual differences as prediction errors. We developed InferAging, a framework that estimates biological age by explicitly modeling individual deviations from chronological age. In progeroid zebrafish (*klotho* mutant; *kl*^−/−^), InferAging identified accelerated and delayed agers within the same chronological age group. A non-invasive variant using behavioral and morphological snapshots reproduced the transcriptome-integrated estimates without molecular input or lifelong tracking. The inferred aging state was associated with metabolic decline, intestinal barrier dysfunction, inflammation, and mucosal immune abnormalities beyond chronological age. These results demonstrate that non-invasive phenotypes can reveal molecularly supported individual aging states.

---

Aging does not progress uniformly across individuals. Even among individuals of the same chronological age, substantial differences exist in physiological and molecular states. Such inter-individual variability reflects underlying heterogeneity in aging rates and trajectories(*1, 2*) and represents a fundamental challenge in aging biology. Accurately capturing this heterogeneity is essential not only for quantifying biological aging beyond chronological age but also for identifying individuals at risk of accelerated aging and those who may benefit from targeted interventions. Biological age has been widely adopted as a metric to quantify the biological progression of aging(*3, 4*). Various approaches have been developed to estimate biological age, ranging from simple phenotypic indices, such as metabolic age and frailty-based measures(*5*), to molecular aging clocks based on DNA methylation, transcriptomic, and proteomic data(*6–8*). Aging-related changes in locomotor activity, exploratory behavior, and body structure across diverse model organisms suggest that non-invasive phenotypes may reflect biological aging states(*9–19*). Consistent with this idea, a behavioral clock based on lifelong recordings in African killifish estimated age from non-invasive behavioral data(*20*).

Despite these advances, most aging clocks remain optimized to predict chronological age or lifespan. Consequently, differences in aging among individuals of the same chronological age are often interpreted as prediction errors in chronological-age estimation. Previous studies have shown that larger training datasets improve chronological-age prediction and reduce these errors in aging clocks(*21–23*). This creates a central paradox for quantifying aging heterogeneity: if individual differences in aging are represented by prediction errors, improving a clock’s age prediction makes those differences harder to quantify. In addition, many informative aging clocks require either molecular profiling or longitudinal phenotypic records acquired over much of the lifespan, limiting their scalability. Thus, a key unresolved problem is how to estimate individual-specific aging deviations as biologically meaningful signals rather than prediction errors. A scalable solution would require non-invasive phenotypic snapshots that do not rely on lifelong tracking, together with molecular evidence supporting the inferred aging states.

Here, we sought to determine whether individual-specific aging deviations supported by internal molecular states can be inferred from single-time-point non-invasive phenotypes. To address this problem, we established InferAging, a hierarchical Bayesian framework that estimates biological age by explicitly modeling each individual’s deviation from chronological age in a progeroid zebrafish model (*kl*^−/−^) (Fig. 1).

**Fig. 1.**
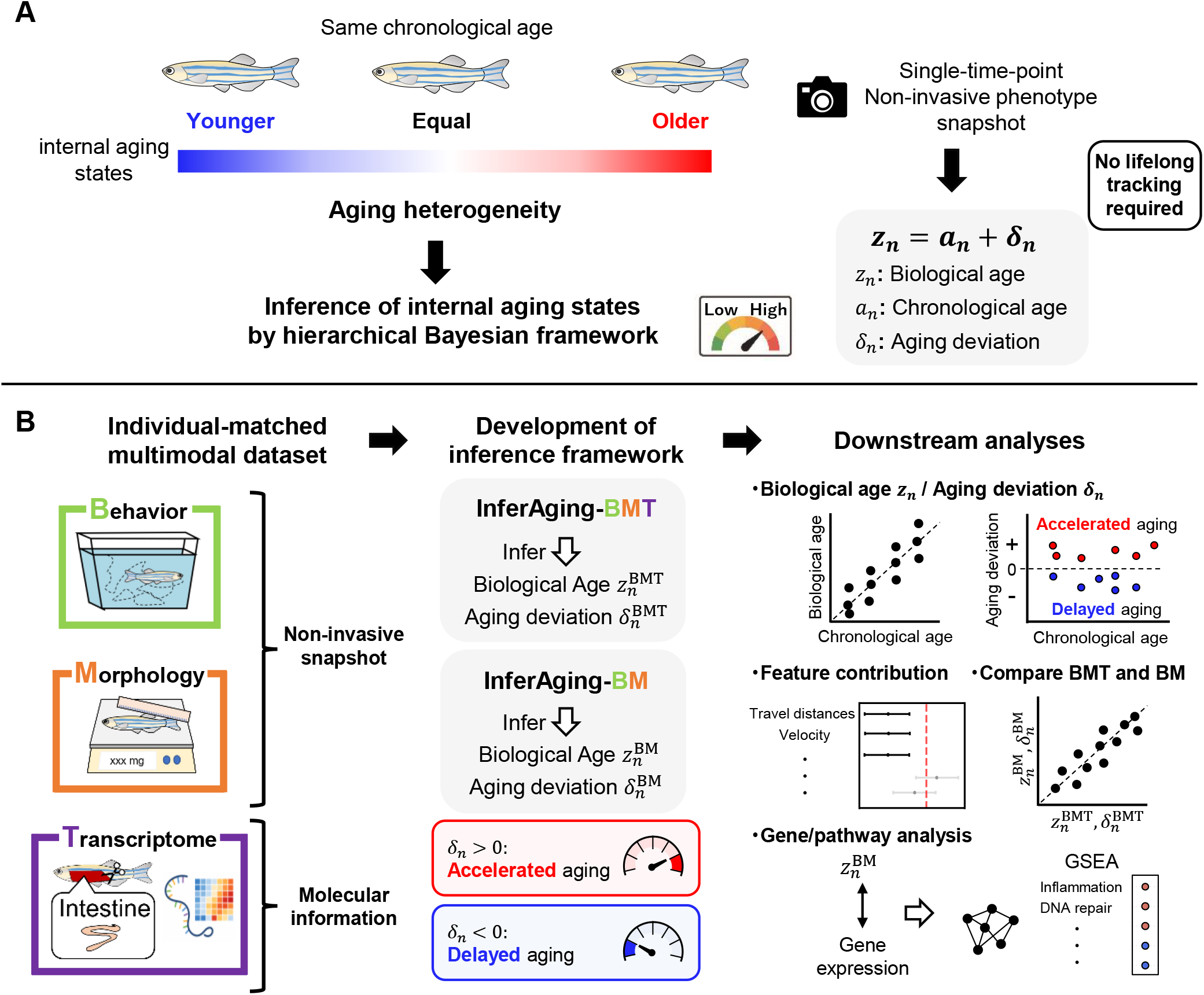
Hierarchical Bayesian framework for modeling inter-individual differences in aging and estimating biological age from non-invasive phenotypic data. (**A**) Conceptual overview of this study. Individuals with the same chronological age can exhibit different internal aging states, reflecting inter-individual heterogeneity in aging. In this framework, biological age (*z*_*n*_) is represented as chronological age (*a*_*n*_) plus an individual-specific aging deviation (*δ*_*n*_), enabling inference of biological age and deviation from chronological age from observable phenotypic data. (**B**) Schematic diagram of analysis flow using InferAging-BMT and InferAging-BM Individual-matched behavioral, morphological, and intestinal transcriptomic data were used to develop InferAging-BMT, whereas InferAging-BM was constructed to estimate biological age and aging deviation using only non-invasive behavioral and morphological data. Downstream analyses included feature contribution analysis, comparison between InferAging-BMT and InferAging-BM estimates, and transcriptome association/pathway analysis to evaluate the biological relevance of InferAging-BM-derived aging estimates.

## Construction of an individual-matched multimodal dataset in progeroid zebrafish

To estimate individual biological age and deviation from chronological age, and to enable downstream analyses of non-invasive age estimation, feature contributions, and molecular programs associated with aging, we constructed an individual-matched multimodal dataset in a vertebrate model system. We used *kl*^−/−^ zebrafish, a progeroid line with a shortened lifespan(*24*), as described in the Materials and Methods, and analyzed individuals at 4, 5, 6, 7, and 8 months post-fertilization (mpf) (n = 21), covering stages from young to old age. For each individual, we obtained non-invasive phenotypic data, including behavioral and morphological measurements, followed by invasive molecular data in the form of intestinal transcriptomic profiles. This sampling scheme enabled direct matching of organism-level phenotypes with internal molecular aging states within the same individuals (fig. S1).

## Continuous and heterogeneous aging obscured by age-group comparisons

We first examined whether conventional age-group comparisons were sufficient to capture aging-related variation in this dataset. Behavioral features such as total distance, velocity, and variance of velocity, as well as morphological traits including body weight, standard length, and total length, generally declined with chronological age (fig. S2 and S3). However, although several behavioral features showed significant overall age effects, pairwise differences were mainly detected between widely separated age groups, whereas adjacent groups exhibited substantial overlap (table S1). Morphological traits showed similar age-dependent trends, but large inter-individual variation limited the detection of statistically significant differences.

We next asked whether molecular data showed a similar pattern. Intestinal transcriptomic profiling revealed progressive age-associated changes, with the number of differentially expressed genes relative to the youngest group increasing with age (fig. S4 and S5). In contrast, comparisons between adjacent age groups identified only limited differences, suggesting that short-term molecular changes were also masked by inter-individual variability within the same age group.

To further assess whether the dataset captured aging as a continuous process, we performed dimensionality reduction using UMAP for each data modality—behavior, morphology, and transcriptome—and visualized the global structure of the data. Across all modalities, samples did not form discrete clusters by age group; instead, they were arranged in chronological order (4–8 mpf) with partial overlap between adjacent groups (Fig. 2, A to C). These results indicate that aging is represented as continuous progression rather than discrete group-wise differences.

**Fig. 2.**
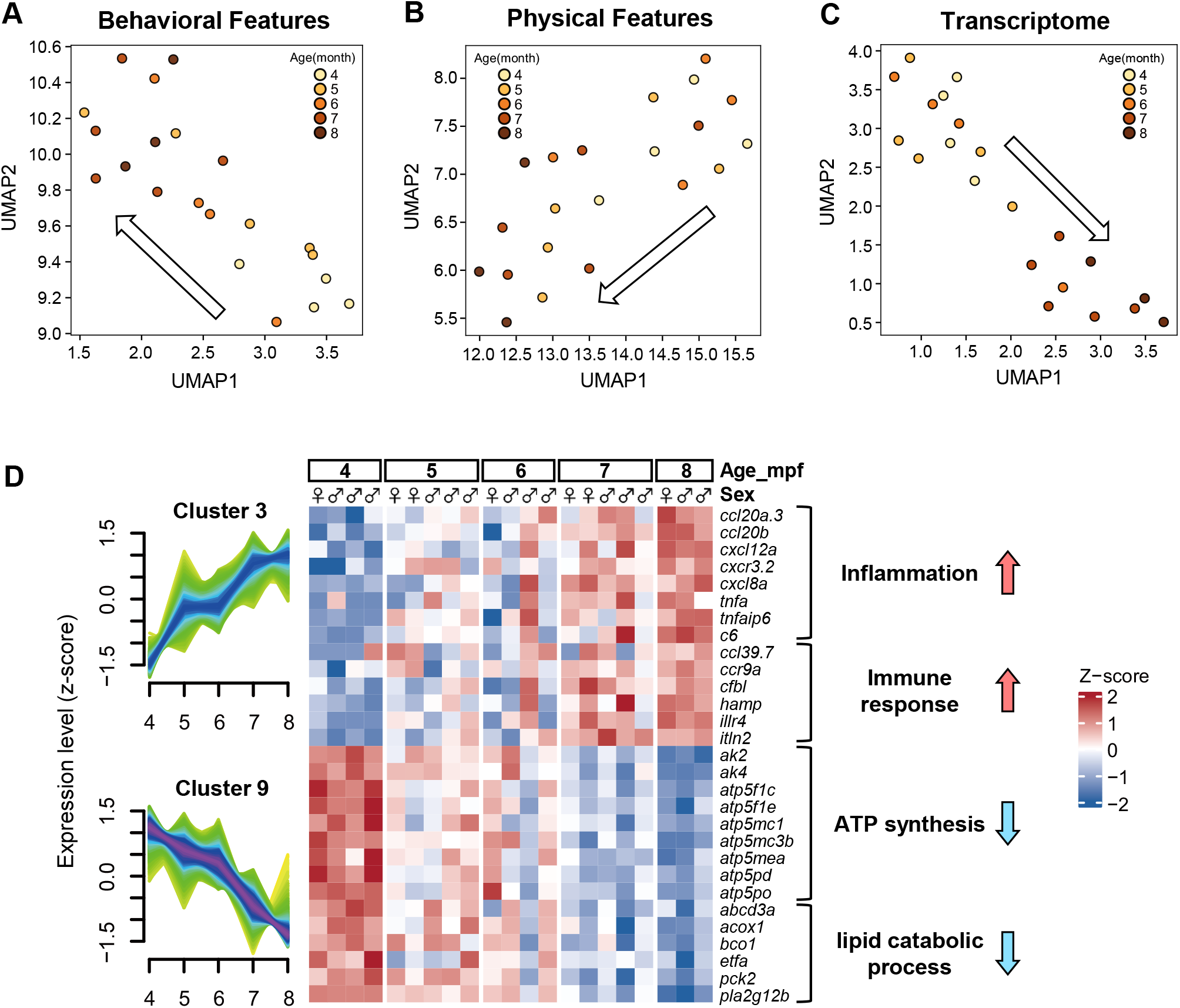
Multimodal analysis of aging phenotypes in *kl*^−/−^ zebrafish. (**A** to **C**) UMAP plots showing the distribution of behavioral (A), morphological (B), and transcriptomic (C) features. Each point represents an individual and is colored according to discrete chronological age groups (light orange: 4 mpf to dark brown: 8 mpf). (**D**) Time-series clustering analysis of transcriptomic data using Mfuzz. Left: standardized (z-scored) expression profiles of representative gene clusters. The x-axis indicates age in months post-fertilization (mpf), and the y-axis indicates standardized expression levels. Right: heatmap showing the expression levels of the identified clusters and representative GO terms revealed by over-representation analysis (ORA). The color scale represents z-scored expression levels (blue: −2, white: 0, red: 2). The color bars above the heatmap indicate chronological age groups (4–8 mpf) and sex (female: red; male: blue). The significance of ORA was assessed by a hypergeometric test with Benjamini–Hochberg correction (q-value < 0.05).

We next evaluated whether the transcriptomic data appropriately reflected biologically meaningful aging states. To uncover temporal patterns masked by inter-individual variability, we performed time-ordering analysis using Mfuzz clustering across all samples. This analysis identified gene clusters exhibiting characteristic age-associated expression patterns (fig. S6). Over-representation analysis revealed that clusters with increasing expression were enriched for inflammatory and immune response pathways, whereas clusters with decreasing expression were enriched for ATP production and lipid metabolism (Fig. 2D and fig. S7). KEGG analysis further confirmed downregulation of metabolic pathways, including oxidative phosphorylation and the citrate cycle (fig. S7B). These features correspond to well-established hallmarks of aging, including inflammaging and mitochondrial dysfunction(*25*).

Taken together, both macroscopic phenotypes and molecular profiles exhibited gradual age-associated shifts while retaining substantial overlap among individuals of the same age. These results indicate that aging in this dataset is better described as a continuous and heterogeneous process rather than discrete age categories. This observation motivated the development of a latent-variable model to infer individual biological age and deviations from chronological age by integrating non-invasive phenotypes with molecular aging readouts.

## Biological age and individual aging deviation estimated by InferAging-BMT

Behavioral, morphological, and transcriptomic data exhibited continuous age-associated changes while retaining inter-individual variability (Fig. 2, A to C). To infer latent aging status from these multivariate data and capture inter-individual heterogeneity, we developed a hierarchical Bayesian model, termed InferAging-BMT (Behavior, Morphology and Transcriptome) (Fig. 3A). In this model, biological age is represented as a latent variable *z*_*n*_ for each individual *n* ∈ {1, … , *N*}. The core assumption of this model is that biological age can be decomposed into chronological age and an individual-specific deviation:

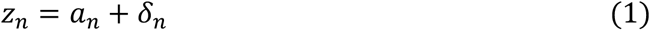

**Fig. 3.**
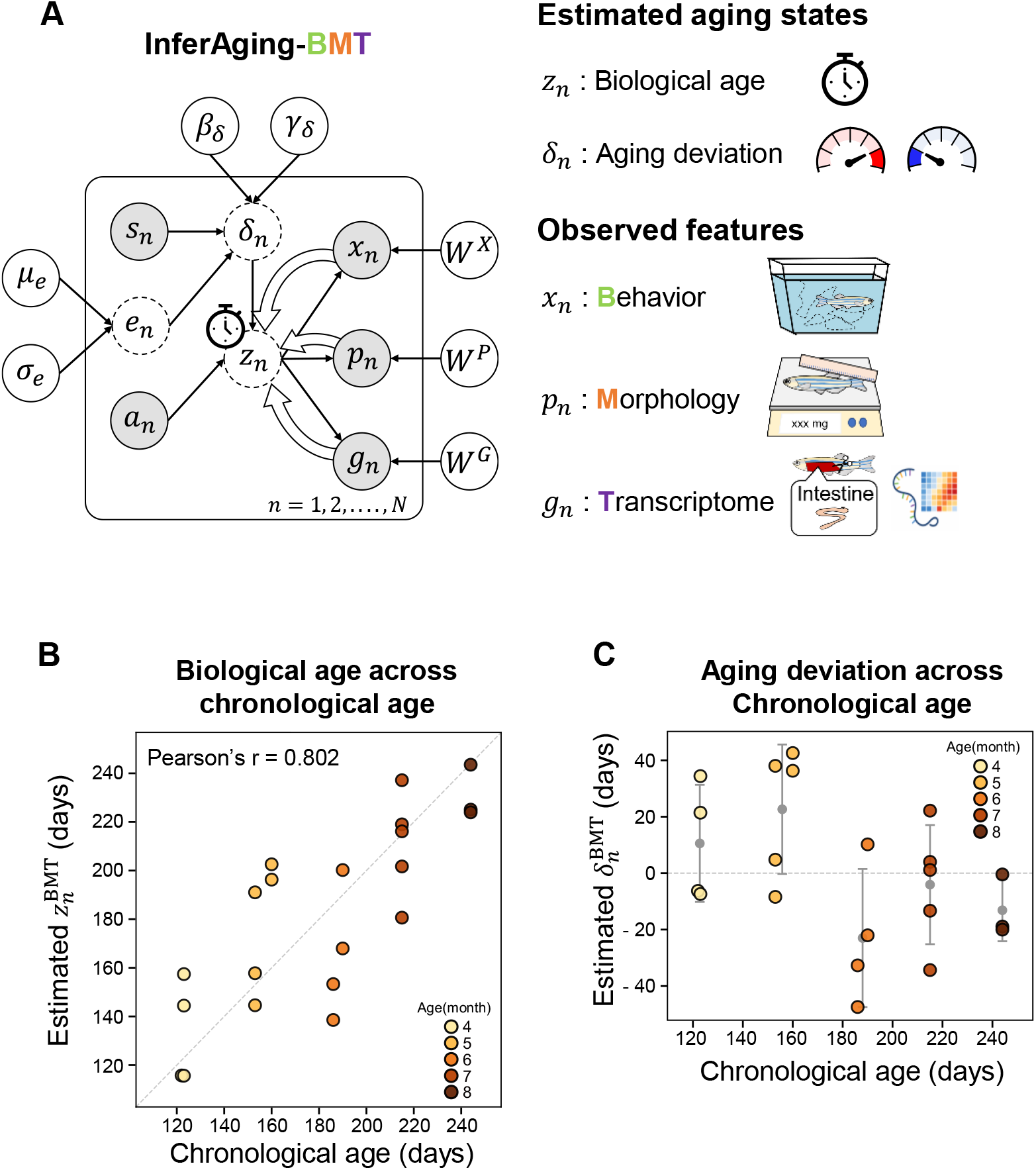
Estimation of biological age using InferAging-BMT. (**A**) Graphical model of InferAging-BMT. Gray nodes indicate observed variables, and white nodes indicate latent variables inferred by the model. *µ*_*e*_, *σ*_*e*_, *β*_*δ*_, *γ*_*δ*_, and *W* (weights for each modality) represent global parameters. For clarity, bias terms, observation noise variances, and some hyperparameters (e.g., *τ*) are omitted from the graphical model. (**B**) Scatter plot of chronological age (*a*_*n*_) and biological age (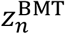) inferred using all observed variables (*x*_*n*_, *p*_*n*_, *g*_*n*_). Pearson’s correlation coefficient (*r*) is shown. (**C**) Distribution of the aging deviation term (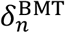) at each age. Each point represents an individual (n = 21), and error bars indicate the standard deviation (s.d.). In (B), the significance of the correlation was tested by Pearson’s correlation analysis.

Here, *z*_*n*_ represents the latent biological age of individual *n, a*_*n*_ is chronological age, and *δ*_*n*_ captures how much the biological age of an individual differs from chronological age (see Materials and Methods). We modeled *δ*_*n*_ as a normally distributed variable whose mean is a linear combination of sex *s*_*n*_, encoded as a binary variable with female = 0 and male = 1, and an unobserved environmental factor *e*_*n*_. Individuals with positive values of *δ*_*n*_ (*δ*_*n*_ > 0) were interpreted as being in a state of accelerated aging relative to their chronological age, whereas individuals with negative values of *δ*_*n*_ (*δ*_*n*_ < 0) were interpreted as being in a state of delayed aging.

The observed data from multiple modalities were then connected to the shared latent aging state *z*_*n*_ through modality-specific observation models. Specifically, observed behavioral data 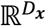, morphological data 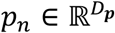, and transcriptomic data 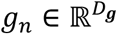 were modeled as linear transformations of *z*_*n*_ with additive Gaussian noise (see Materials and Methods). This model structure represents a biological process in which all observed modalities change coordinately, driven by the shared latent aging state *z*_*n*_.

To assess whether this formulation captures inter-individual differences in aging, we integrated behavioral data *x*_*n*_ , morphological data *p*_*n*_ , and transcriptomic data *g*_*n*_ , and performed Bayesian inference to estimate the posterior distributions of model parameters and individual biological age. After fitting InferAging-BMT, we denote the posterior estimate of *z*_*n*_ as 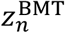, and the corresponding aging deviation as 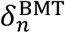 . Posterior predictive checks (PPCs) indicated that the model accurately reproduced the observed data distributions, demonstrating good model fit (fig. S8). The estimated 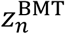 showed a positive correlation with chronological age a_n (n = 21, Pearson’s r = 0.802, p-value = 1.22×10^-5^, Fig. 3B and Table 1). Notably, 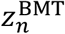 did not simply follow *a*_*n*_, but captured variability among individuals of the same chronological age. Indeed, analysis of the distribution of 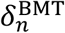 showed that, even within the same 4–7 mpf age groups, individuals with accelerated aging (*δ*_*n*_ > 0) and those with delayed aging (*δ*_*n*_ < 0) were clearly distinguished (Fig. 3C). These results indicate that InferAging-BMT captures not only chronological age but also quantitative inter-individual differences in aging.

**Table 1.** Comparison of biological age prediction performance across aging estimation models.

| Model | Input Modalities | Estimated biological age | Evaluated against | Pearson's $r$ | $R^2$ | MAE (Days) |
| --- | --- | --- | --- | --- | --- | --- |
| InferAging-BMT | $X, P, G$ | $z_n^{\text{BMT}}$ | Chronological age | 0.802 | — | — |
| InferAging-BMT (5fold-CV) | $X, P, G$<br>($G$: Use only during training) | Predicted $\hat{z}_n^{\text{BMT}}$ | $z_n^{\text{BMT}}$ | 0.988 | 0.976 | 8.41 |
| InferAging-BM | $X, P$ | $z_n^{\text{BM}}$ | Chronological age | 0.694 | — | — |
| InferAging-BM | $X, P$ | $z_n^{\text{BM}}$ | $z_n^{\text{BMT}}$ | 0.981 | 0.961 | 5.78 |

## Non-invasive prediction of transcriptome-integrated biological age by masked-transcriptome cross-validation

Next, we tested whether the internal aging state could be predicted using only non-invasive data (*x*_*n*_, *p*_*n*_) without requiring tissue collection or dissection. To this end, we treated 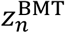 inferred by InferAging-BMT from the full dataset including transcriptomic data as the reference biological age in this study, and evaluated the predictability of 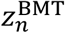 using only non-invasive data. Predictive accuracy was evaluated using repeated 5-fold cross-validation. During training, InferAging-BMT parameters were estimated using all observed data of training samples (*X*_train_, *P*_train_, *G*_train_), enabling the model to learn the latent relationships among behavior, morphology, and transcriptome. During testing, by contrast, invasive transcriptomic data *G*_test_ were masked, and only non-invasive behavioral and morphological data of test samples (*X*_test_, *P*_test_) were used as input. Under this condition, we predicted 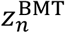 from non-invasive data alone by fixing the learned parameters and performing Bayesian inference of the latent variables for each individual (Fig. 4A). To reduce the influence of random data partitioning and ensure robustness, this 5-fold cross-validation procedure was repeated 100 times with different fold assignments, and the final performance metrics were calculated by averaging the results across these repeated trials.

**Fig. 4.**
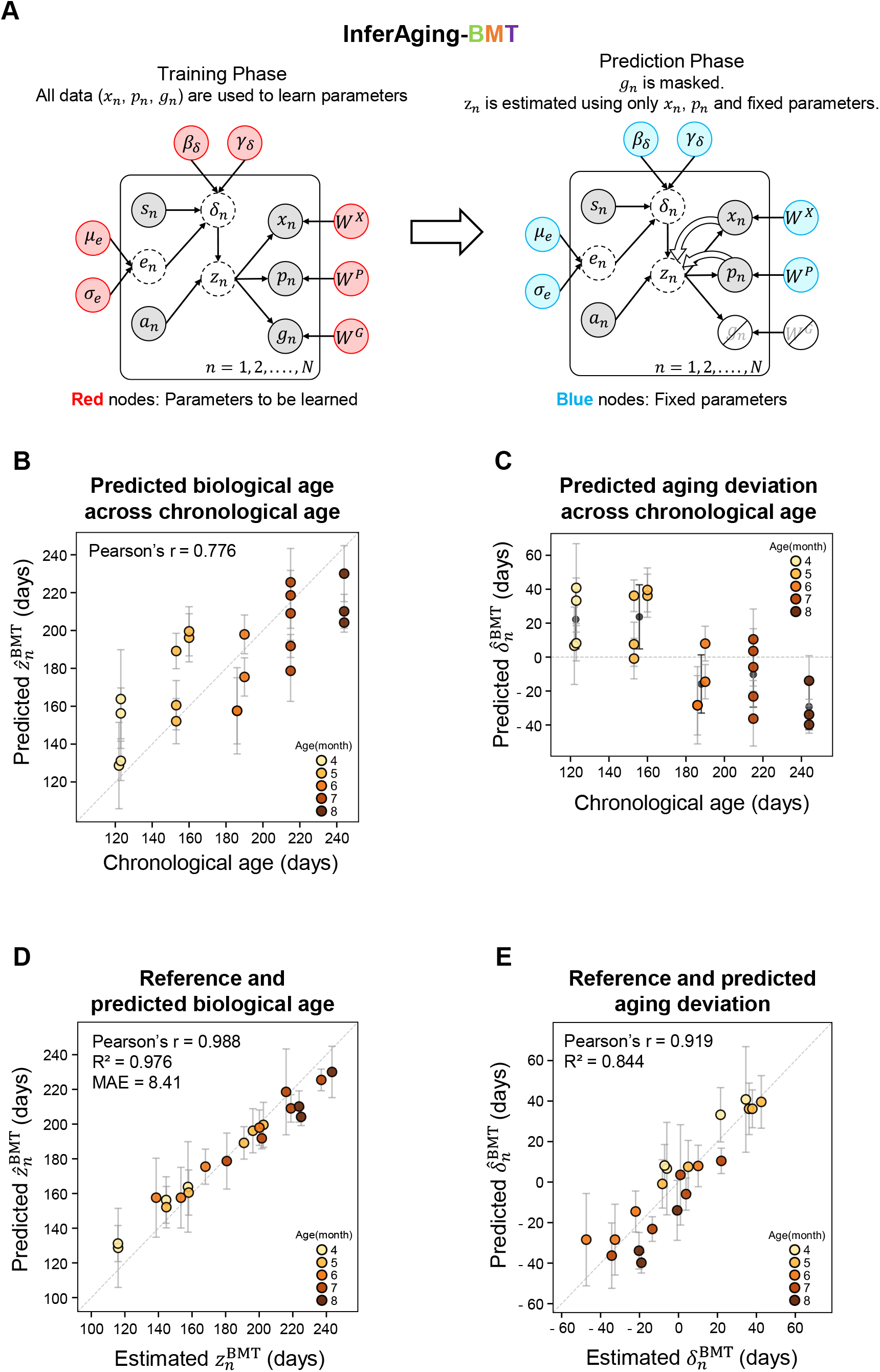
Non-invasive prediction and validation of biological age using InferAging-BMT with repeated 5-fold cross-validation. (**A**) Graphical model of the non-invasive aging estimation framework using repeated 5-fold cross-validation. In the test phase, transcriptomic data (*g*_*n*_) were masked, and biological age (predicted 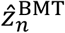) was estimated using only non-invasive behavioral (*x*_*n*_) and morphological (*p*_*n*_) features. (**B**) Scatter plot of chronological age (*a*_*n*_) and biological age predicted from non-invasive data only (predicted 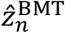). Pearson’s correlation coefficient (*r*) is shown. (**C**) Distribution of the predicted aging deviation term (predicted 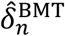) at each age. Each point represents an individual (n = 21). Gray error bars indicate the standard deviation (s.d.) for each individual across 100 repeated 5-fold cross-validation trials, and black error bars indicate the standard deviation among individuals within each age group. (**D**) Scatter plot of the reference biological age estimated from the full dataset (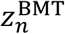) and the biological age predicted for each individual using only non-invasive data (predicted 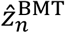). Pearson’s correlation coefficient (*r*), coefficient of determination (*R*^2^), and mean absolute error (MAE) are shown. (**E**) Scatter plot of the reference aging deviation term (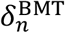) and the deviation term predicted from non-invasive data (predicted 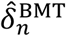). Pearson’s correlation coefficient (*r*) and coefficient of determination (*R*^2^) are shown. In (B), (D), and (E), each point represents an individual (n = 21), and error bars indicate the standard deviation (s.d.) across 100 repeated 5-fold cross-validation trials. The dotted line in these panels indicates the identity line (y = x), and the significance of correlations was tested by Pearson’s correlation analysis. In (C), the dotted line indicates 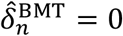= 0.

The predicted 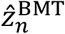 based only on non-invasive data remained positively correlated with chronological age *a*_*n*_ (n = 21, Pearson’s r = 0.776, p-value = 3.57×10^-5^, Fig. 4B). Importantly, even in the absence of transcriptomic input during testing, inter-individual variability was preserved among individuals of the same age. Consistently, the distribution of predicted 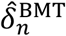 showed that individuals with accelerated 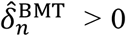 and delayed 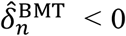 aging coexisted within the 5–7 mpf groups (Fig. 4C). These results suggest that behavioral and morphological data alone are sufficient to capture inter-individual differences in aging, even without transcriptomic information at the prediction phase.

We next evaluated the accuracy of biological age prediction using only non-invasive data by comparing 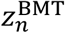 inferred using full data with 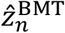 predicted from only non-invasive data.

Predicted 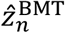 showed a strong correlation with 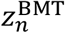 (n = 21, Pearson’s r = 0.988, *R*^2^ = 0.976, p-value = 7.57×10^-17^, MAE = 8.41 days; Fig. 4D). Similarly, predicted 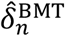 was significantly correlated with 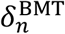 (Pearson’s r = 0.919, *R*^2^ = 0.844, p = 4.26 × 10^−9^; Fig. 4E and Table 1), indicating that inter-individual differences in aging progression were preserved. These results demonstrate that, once trained with the full multimodal dataset including transcriptomic data, InferAging-BMT can accurately reproduce reference biological age and aging-deviation estimates from non-invasive phenotypic inputs alone.

## Feature contributions to biological age estimation

To identify features contributing to biological age estimation, we analyzed the posterior distributions of weight coefficients *W* inferred by InferAging-BMT (Fig. 5A and table S2). In this model, *W* reflects the direction and magnitude of the association between each feature and biological age *z*_*n*_: positive values indicate that the feature is associated with higher *z*_*n*_, whereas negative values indicate that it is associated with lower *z*_*n*_ . Features whose 95% credible intervals (CIs) excluded zero were considered to contribute to aging estimation. Among behavioral features (*X*), six activity-related features contributed to aging estimation and showed negative weights *W*^*X*^ (Fig. 5B). These included mean swimming velocity, total swimming distance, and frequency of entry into the upper tank region, all of which had 95% CIs below zero (e.g., Velocity: mean = −0.76, 95% CI [−1.25, −0.33]). Among morphological features (*P*), ten of twelve features—including total length, body weight, and body depth—also showed negative weights *W*^*P*^ (Fig. 5C).

**Fig. 5.**
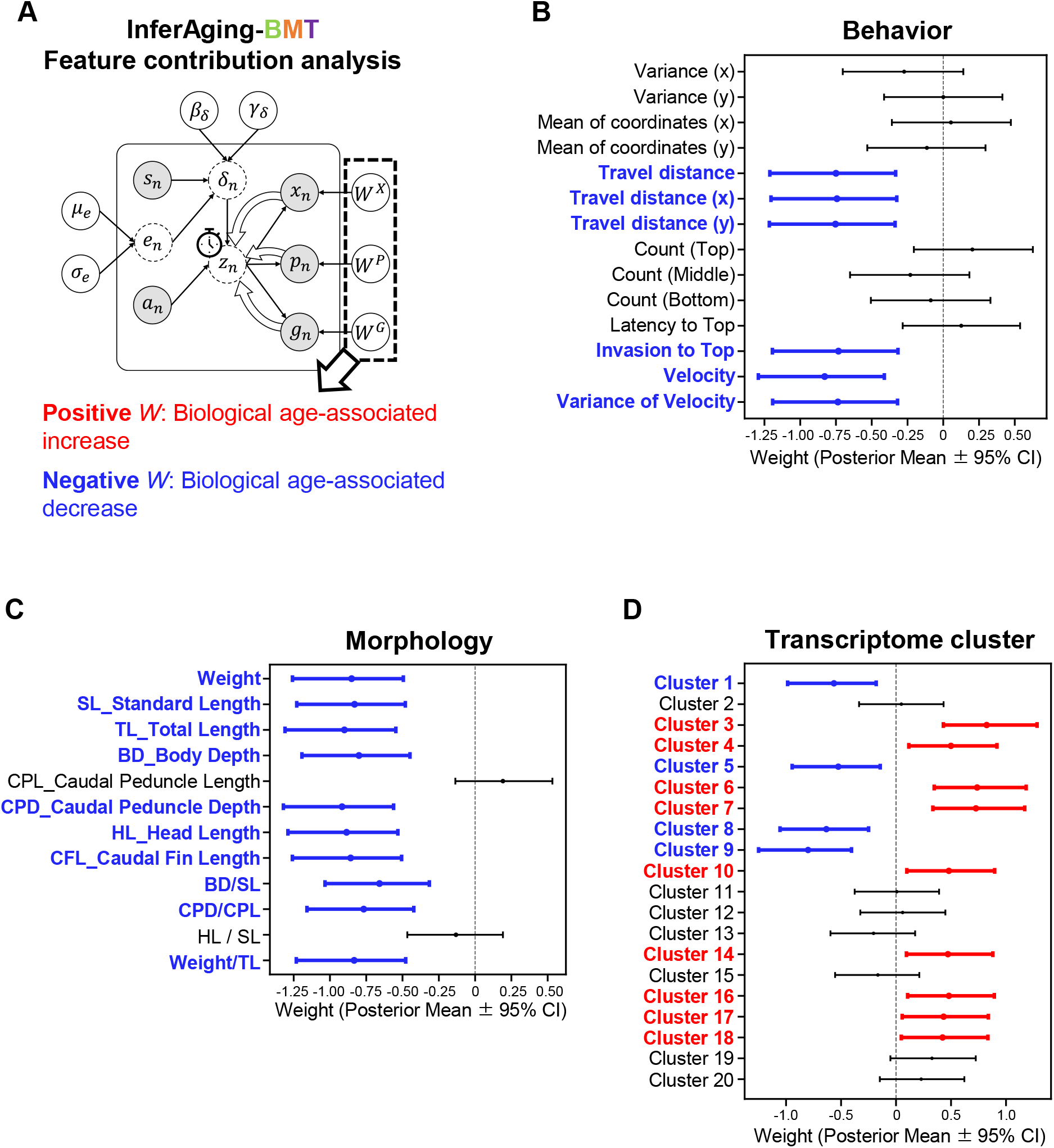
Feature contribution analysis based on posterior weights estimated by InferAging-BMT. (**A**) Schematic overview of the feature contribution analysis. Positive and negative weights represent biological age-associated increases and decreases, respectively. (**B**) Forest plot of the posterior weight coefficients (*W*^*X*^) for behavioral features. (**C**) Forest plot of the posterior weight coefficients (*W*^*P*^) for morphological features. (**D**) Forest plot of the posterior weight coefficients (*W*^*G*^) for transcriptome cluster features. In (B), (C), and (D), each point indicates the posterior mean, and the horizontal error bars indicate the 95% credible interval (CI), calculated as the 2.5th to 97.5th percentiles of the posterior distribution. Red points and error bars indicate positive weights whose 95% CI excludes 0, blue points and error bars indicate negative weights whose 95% CI excludes 0, and black points and error bars indicate weights whose 95% CI includes 0. The vertical dashed line indicates the position at which the weight coefficient equals 0.

For transcriptomic features (*G*), we analyzed weight coefficients *W*^*G*^ for 20 gene clusters identified by Mfuzz (Fig. 5D). Thirteen clusters showed significant contributions (95% CI excluding zero), including clusters with positive weights (Clusters 3, 4, 6, 7, 10, 14, 16, 17, 18) and clusters with negative weights (Clusters 1, 5, 8, 9). These clusters were consistent with enrichment results from earlier analyses (Fig. 2D and fig. S7). For example, Cluster 3 (positive weight) was enriched for GO terms related to inflammatory and immune responses, whereas Cluster 9 (negative weight) was enriched for ATP synthesis and lipid metabolism (Table 2 and fig. S7). Notably, several clusters (e.g., Clusters 4 and 17) made significant contributions despite showing no significant ORA enrichment (Table 2 and fig. S7). Gene-level inspection revealed the presence of key aging-related genes, including *rptor* and *fgf21* in Cluster 4, and *prkaa1* and *kmt5b* in Cluster 17. These findings suggest that InferAging-BMT capture aging-relevant signals not fully summarized by existing functional annotations.

**Table 2.** Biological characteristics and key genes of representative gene expression clusters contributing to biological age estimation.

| Cluster | Mean weight [95% CI] | Expression Trend | Representative GO terms / pathways | Representative genes |
| --- | --- | --- | --- | --- |
| 3 | 0.824 [0.430, 1.282] | Upregulated | Inflammatory response<br>Immune response | *illr4, tnfa* |
| 9 | - 0.800 [- 1.250, - 0.405] | Downregulated | ATP synthesis<br>lipid catabolic process | *atp5f1c, acox1* |
| 4 | 0.499 [0.116, 0.918] | Fluctuating | No significant enrichment | *rptor, adipor1a* |
| 17 | 0.432 [0.056, 0.839] | Fluctuating | No significant enrichment | *prkaa1, kmt5b* |

## Non-invasive estimation of biological age and individual aging deviation by InferAging-BM

The analysis based on InferAging-BMT demonstrated that biological age *z*_*n*_ can be predicted without transcriptomic input during testing. Furthermore, posterior analyses revealed that behavioral and morphological features contribute substantially to this estimation. Motivated by these findings, we developed InferAging-BM (Behavior and Morphology), a fully non-invasive model that estimates latent biological age using only behavioral and morphological data (*X, P*), without incorporating transcriptomic data (*G*) during training. Specifically, we removed the transcriptomic observation model and its associated parameters, including the weight coefficients *W*^*G*^ , from InferAging-BMT. The resulting model was then applied to infer the posterior distribution of the latent variable *z*_*n*_ for all individuals based solely on behavioral and morphological observations (Fig. 6A).

**Fig. 6.**
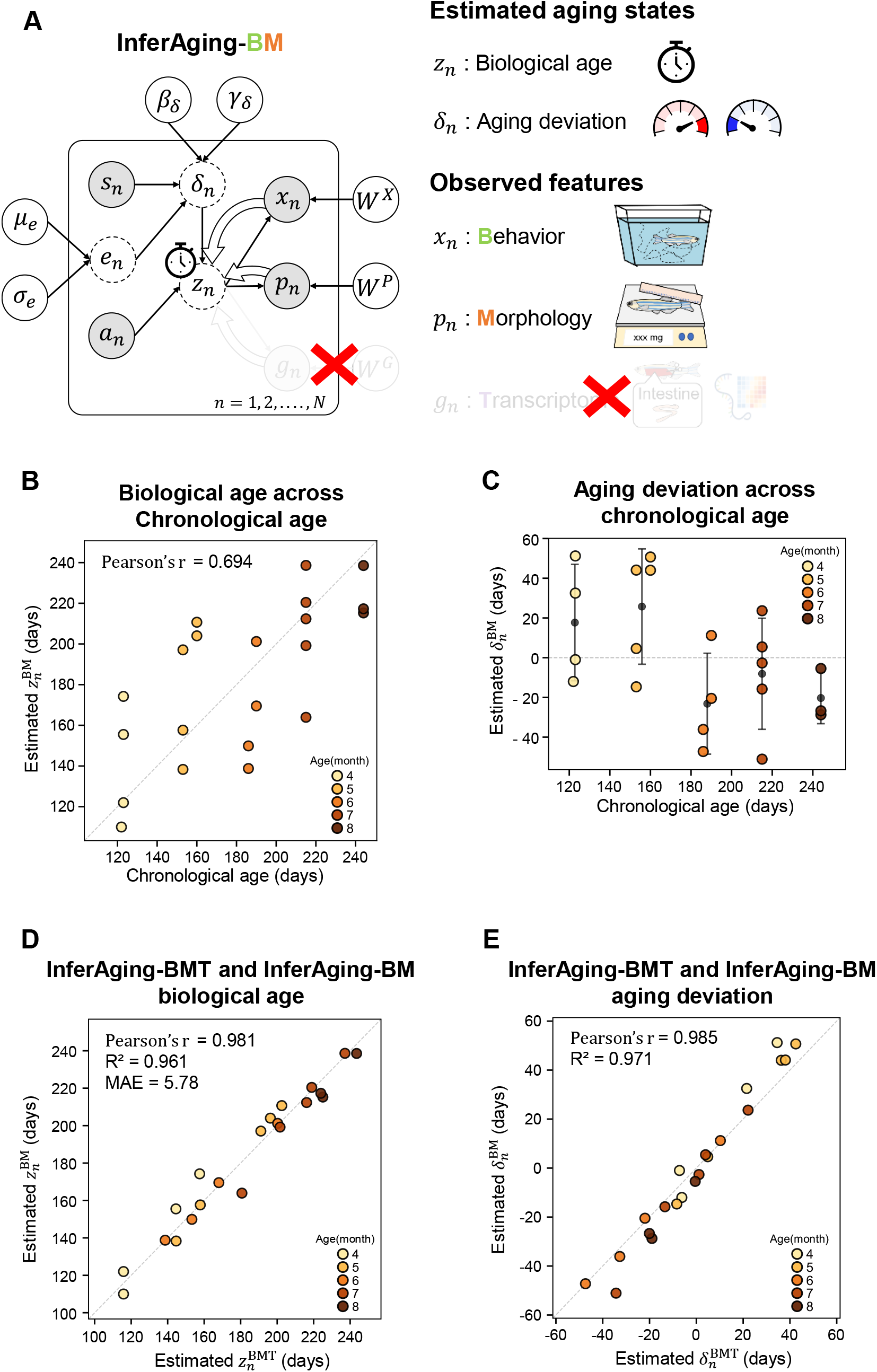
Reconstruction of biological age by the transcriptome-independent, fully non-invasive model InferAging-BM. (**A**) Graphical model of InferAging-BM. (**B**) Scatter plot of chronological age (*a*_*n*_) and biological age estimated by InferAging-BM (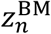). Pearson’s correlation coefficient (*r*) is shown. (**C**) Distribution of the aging deviation term estimated by InferAging-BM (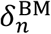) at each age. (**D**) Scatter plot of 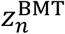 estimated using the full dataset and 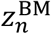. Pearson’s correlation coefficient (*r*), coefficient of determination (*R*^2^), and mean absolute error (MAE) are shown. (E) Scatter plot of 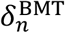 estimated using the full dataset and 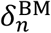 estimated by InferAging-BM. Pearson’s correlation coefficient (*r*) and coefficient of determination (*R*^2^) are shown. In (B), (D), and (E), the dotted line indicates the identity line (y = x). Each point represents an individual (n = 21). In (B), (D), and (E), the significance of the correlation was tested by Pearson’s correlation analysis. In (C), the dotted line indicates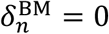.

Biological age estimated by InferAging-BM, 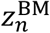 , showed a significant positive correlation with chronological age (Pearson’s r = 0.694, p-value = 4.85 × 10^−4^; Fig. 6B). Consistent with the InferAging-BMT results, the distribution of 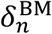 indicated the coexistence of individuals with accelerated (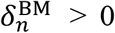) and delayed (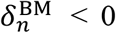) aging within the same age groups (4–7 mpf; Fig. 6C). Notably, 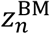 showed remarkably high agreement with 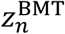 (Pearson’s r = 0.981, *R*^2^ = 0.961, p = 6.82 × 10^−15^, MAE = 5.78 days; Fig. 6D and Table 1). Similarly, 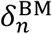 was highly consistent with 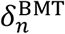 (Pearson’s r = 0.985, *R*^2^ = 0.971, p-value = 4.66 × 10^−16^; Fig. 6E). These results suggest that behavioral and morphological phenotypes contain sufficient information to reproduce transcriptome-integrated biological age estimates without transcriptomic input. Thus, within the scope of this dataset, biological age reflecting molecular aging states can be accurately estimated from non-invasive phenotypic inputs alone.

To assess the robustness of InferAging-BM estimates to variation in the composition of chronological age groups, we performed two additional sensitivity analyses. First, in a one-age-group-dropout analysis, InferAging-BM was refitted after excluding one chronological age group at a time, and estimates for the remaining shared individuals were compared with those obtained using the full dataset (fig. S9A). Across all dropout conditions, aging deviation 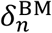 estimates remained highly consistent with full-data InferAging-BM estimates (*R*^2^ = 0.910– 0.987, MAE = 2.94–8.07 days, sign consistency = 0.813–1.000; fig. S9, B and C). Biological age 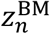 estimates also showed close alignment with full-data estimates, supporting the stability of InferAging-BM-derived 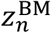 under age-group dropout (fig. S9D). Second, in an age-imbalance resampling analysis, InferAging-BM was refitted using randomly imbalanced subsets while retaining all chronological age groups, and estimates for individuals included in each subset were compared with those obtained using the full dataset (fig. S10A). Across 30 resampling trials, aging deviation 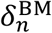 estimates remained generally consistent with full-data InferAging-BM estimates (*R*^2^ = 0.799–0.994, MAE = 1.84–11.38 days, sign consistency = 0.733–1.000; fig. S10, B and C). Biological age 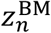 estimates also showed close agreement with full-data estimates (fig. S10D). Together, these analyses suggest that InferAging-BM-derived aging deviation and biological age estimates were not strongly dependent on the relative composition of chronological age groups.

## Molecular aging states reflected by non-invasive biological age

To assess whether InferAging-BM captures biologically meaningful aging states, we evaluated the relationship between the estimated 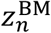 and transcriptomic data that were not used during model training. Spearman’s rank correlation analysis identified 412 genes significantly associated with 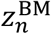 (219 positively and 193 negatively correlated genes; q < 0.10; Fig. 7A), suggesting that InferAging-BM-derived biological age is linked to specific gene expression patterns.

**Fig. 7.**
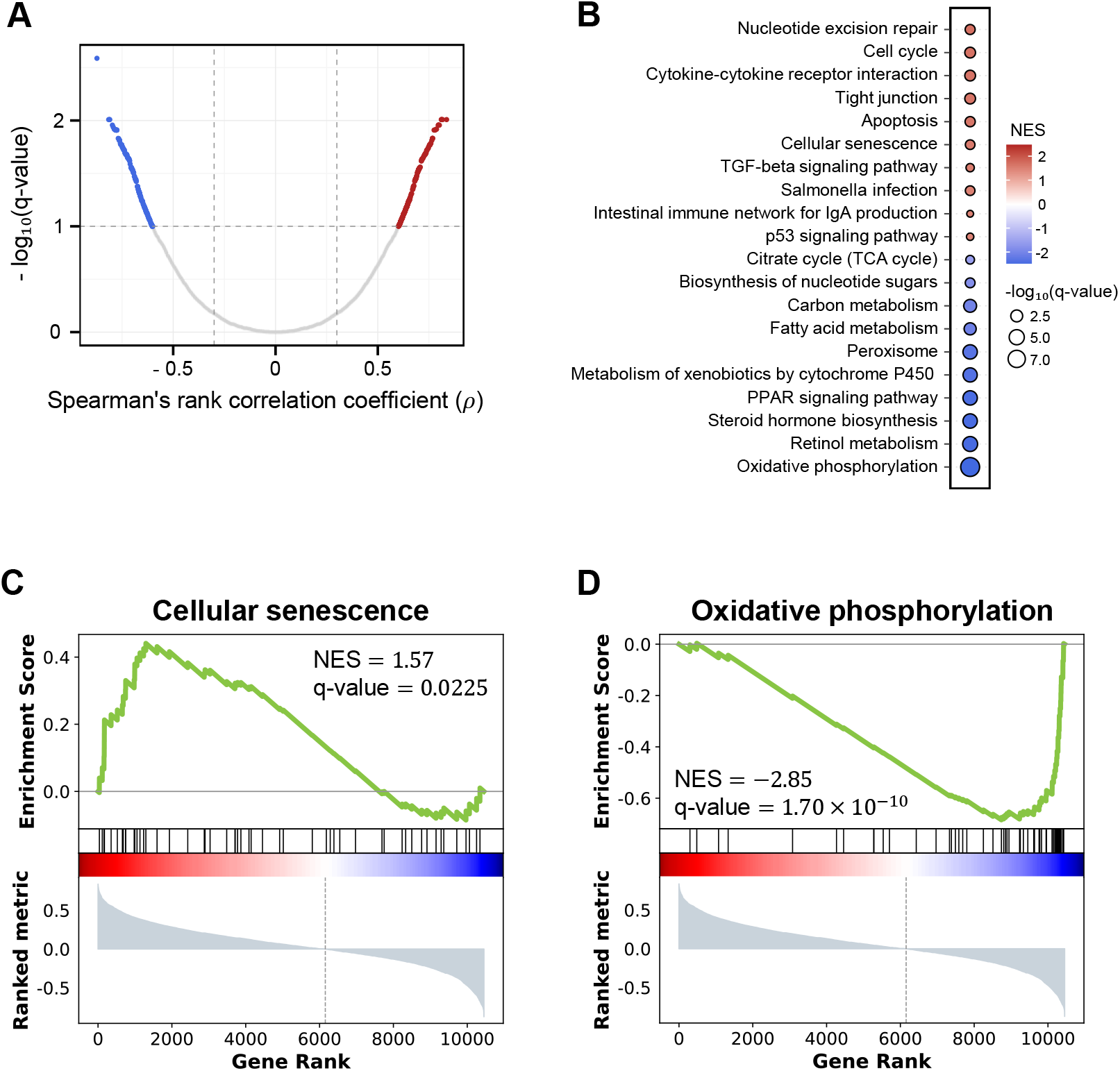
Gene expression profiles and aging-related pathways significantly correlated with 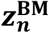. (**A**) Volcano plot showing the correlation between the expression level of each gene and biological age estimated by InferAging-BM (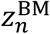). The x-axis represents Spearman’s rank correlation coefficient, and the y-axis represents −log_10_(q-value). The black dashed line indicates the threshold of q-value = 0.10. Among genes satisfying q < 0.10, positively correlated genes are shown in red and negatively correlated genes are shown in blue. (**B**) Dot plot of representative pathways identified by GSEA of KEGG pathways using the Spearman’s rank correlation coefficients for all genes. The color bar indicates the normalized enrichment score (NES). Dot size represents −log_10_(q-value). (**C**) GSEA enrichment plot for “Cellular senescence.” (**D**) GSEA enrichment plot for “Oxidative phosphorylation.”

To characterize the underlying biological processes, we performed Gene Set Enrichment Analysis (GSEA) using a correlation-ranked gene list. KEGG pathway analysis revealed significant enrichment of multiple aging-related pathways (Fig. 7B and fig. S11). Genes positively correlated with 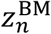 were enriched for inflammatory and cellular stress response pathways, including “Cytokine–cytokine receptor interaction” (NES = 1.64), “Cellular senescence” (NES = 1.57; Fig. 7C), “Apoptosis” (NES = 1.62), and “p53 signaling pathway” (NES = 1.46). Pathways related to mucosal immunity and barrier function, such as “Intestinal immune network for IgA production” (NES = 1.49) and “Tight junction” (NES = 1.64), were also enriched. In contrast, genes negatively correlated with 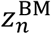 were enriched for mitochondrial and metabolic pathways, including “Oxidative phosphorylation” (NES = −2.85; Fig. 7D), “Citrate cycle” (NES = −1.81), and “Fatty acid metabolism” (NES = −2.07). Lipid metabolism–related pathways such as “PPAR signaling pathway” (NES = −2.43) and “Peroxisome” (NES = −2.29) were also significantly enriched. These results indicate that the biological age estimated by InferAging-BM (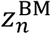) reflects canonical molecular hallmarks of aging, despite being inferred exclusively from non-invasive phenotypic data.

## Discussion

Accurately capturing inter-individual differences in aging among individuals of the same chronological age remains a central challenge in aging research(*26*). Many existing aging clocks prioritize prediction accuracy and consequently treat such differences as noise. Moreover, non-invasive behavioral approaches often require lifelong tracking, limiting their scalability. Here, we address these conceptual and technical limitations by establishing a hierarchical Bayesian framework (InferAging-BMT/InferAging-BM) that explicitly models inter-individual variability in aging as a biologically meaningful signal. In particular, the InferAging-BM approach shifts the existing paradigm, which has relied on lifelong tracking or invasive omics analysis, and demonstrates that biological age supported by internal molecular states can be accurately estimated from single-time-point non-invasive phenotypic snapshots alone.

Many existing aging clocks have been constructed as regression models, such as Elastic Net, that accurately predict chronological age. However, because these approaches are optimized to minimize prediction error relative to chronological age, they introduce a conceptual paradox: as prediction accuracy improves, biologically meaningful inter-individual differences are increasingly suppressed as noise. In this study, we addressed this paradox by constructing the InferAging framework, in which biological age *z*_*n*_ is formulated as two components: a baseline defined by chronological age and an individual-specific deviation from that baseline *δ*_*n*_. This mathematical structure enabled the model to maintain age-estimation capability while naturally extracting inter-individual differences in aging that would otherwise be buried in prediction error as biological signals. Thus, we successfully constructed a model that reflects individual differences in aging.

A particularly important point is the performance and biological significance of InferAging-BM, which uses only non-invasive phenotypes as input. For a model that predicts biological age from behavior and morphology to be truly useful biologically, its estimates must not merely reflect external phenotypic changes, but must accurately reflect internal molecular aging programs. In this study, we directly addressed this requirement by obtaining behavioral, morphological, and intestinal transcriptomic datasets from the same individuals. Although tissue collection was required to construct and validate this dataset, the demonstration using these integrated data showed that InferAging-BM, which does not incorporate transcriptomic information at all during training or inference, produces estimates that are strongly linked to internal molecular aging states. These findings suggest that organism-level phenotypes can serve as compressed readouts of internal molecular aging states.

The biological age 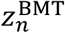 estimated by InferAging-BMT reflects a biologically meaningful state rather than a mere proxy for chronological age. The model assigned strong negative weights to behavioral and morphological features related to activity level and body condition (Fig. 5, B and C), capturing core frailty-associated phenotypes such as reduced activity and emaciation. These features are well-established indicators of declining healthspan in mammalian and human aging(*5, 27, 28*). Concurrently, the model captured key molecular hallmarks of aging, including increased inflammation and reduced ATP synthesis (Fig. 5D and Table 2). Together, these results suggest that phenotypic features such as reduced activity and body condition serve as robust surrogate markers of underlying molecular aging processes.

Notably, some gene clusters that contributed significantly to aging estimation did not show enrichment in standard ORA analyses. However, gene-level inspection revealed that these clusters contained known aging-related genes, including *rptor, fgf21, prkaa1*, and *kmt5b* (Table 2). This suggests that signals overlooked by conventional differential expression analyses can be recovered as biologically meaningful features through integrative modeling of phenotypic and transcriptomic data. Consistent with this, biological age estimated from phenotypes alone (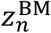) recapitulated molecular-level functional changes, as supported by GSEA (Fig. 7 and fig. S11). Increasing 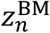 was associated with activation of pathways related to inflammation, cellular senescence, and intestinal homeostasis, including barrier function and mucosal immunity. In contrast, mitochondrial energy metabolism pathways, represented by oxidative phosphorylation, the citrate cycle, and fatty acid metabolism, were markedly suppressed, suggesting mitochondrial dysfunction, reduced ATP production, and disruption of tissue homeostasis(*25*). Importantly, these tissue-specific changes were not fully captured by chronological age-group comparisons, indicating that InferAging-BM-derived biological age reflects not only global aging signals but also local disruptions in tissue homeostasis. These findings suggest that transcriptomic signatures of intestinal dysfunction, traditionally assessed using invasive methods, can be inferred non-invasively and quantitatively from phenotypic snapshots. Furthermore, genes co-expressed within these clusters may include previously unrecognized regulators of aging, warranting future functional validation.

The framework proposed in this study provides a flexible foundation for modeling biological aging and can be further extended in several directions. First, although the present study was based on a relatively small dataset (n = 21), the framework successfully captured key aging-associated signals. Future analyses using larger datasets will enable the development of more expressive models that account for complex gene expression dynamics, including multimodality. Second, while this study relied on cross-sectional data, our results demonstrated that inter-individual differences in aging can be robustly extracted from snapshot observations. Integration with longitudinal data will further allow direct modeling of individual aging trajectories and dynamic responses to interventions. Third, although we used a progeroid model (*klotho* mutant), the framework itself is general and can be applied to natural aging in wild-type individuals and across different tissues. This also provides a basis for extending the model to sex-specific aging analyses. Finally, while experiments in the present study were conducted under controlled single-housing conditions, housing environment and social context may influence aging-related phenotypes. Indeed, juvenile housing density has been shown to affect growth, fecundity, lifespan, and aging-related transcriptomic programs in African turquoise killifish(*29*). Therefore, future validation under more complex environments, including group housing and social interactions, will be important for extending this framework to more physiological settings and for incorporating environmental covariates into the estimation of individual aging trajectories.

In summary, we present a hierarchical Bayesian framework that explicitly captures inter-individual differences in aging and enables accurate estimation of biologically meaningful age from non-invasive phenotypic data. This approach provides a scalable foundation for quantifying aging dynamics across individuals, conditions, and biological systems.

## Materials and Methods

### Experimental animal model

Zebrafish (*Danio rerio*) have a relatively long lifespan of approximately 3–4 years, which has posed a challenge for their application to aging research. However, the *klotho* mutant (*kl*^−/−^), a progeroid mutant, is known to have a shortened lifespan of approximately 9 months. Therefore, in this study, we used *kl*^−/−^ as an experimental animal model in which the progression of aging can be analyzed over a short period(*24*). The *kl*^+/−^ was maintained in system water containing 0.03% salt at 28.5°C under a 14 h light/10 h dark photoperiod. To minimize the influence of genetic background on inter-individual differences in aging, sibling *kl*^−/−^ zebrafish derived from the same *kl*^+/−^ breeding pair were used in all experiments. Fish were fed twice daily according to their developmental stage. Larvae from 5 days post-fertilization (5 dpf) to 1 mpf were fed Hikari Labo 130 (Kyorin). Juveniles from 1 to 3 mpf were fed brine shrimp (Kitamura) and Hikari Labo 130 (Kyorin), and adult fish older than 3 mpf were fed brine shrimp.

### Genotyping

Genomic DNA was extracted from tail fin clips using a standard protocol. Genotyping was performed by allele-specific PCR using the KAPA Taq Extra PCR Kit (Kapa Biosystems) and an Applied Biosystems 2720 Thermal Cycler (Thermo Fisher Scientific). Unlike the previously reported method based on restriction enzyme digestion(*24*), allele-specific PCR was used to separately detect the wild-type and mutant (*kl*^*sa*18644^) alleles. The primers and cycling conditions were as follows. To detect the wild-type allele, the forward primer 5’-CCCGTCCCTTGTTTGTAGATGGAGATTAT-3’ and reverse primer 5’-CTCATTCATCCACATACCTTTTAGCGTCTCC-3’ were used. PCR amplification was performed with an initial denaturation at 94°C for 2 min, followed by 32 cycles of denaturation at 94°C for 10 s, annealing at 68°C for 30 s, and extension at 72°C for 30 s, with a final extension at 72°C for 1 min. To detect the mutant allele, the forward primer 5’-ACTTTCCCAAGAAGAAAAAACATCATAGG-3’ and reverse primer 5’-TTGTCCTTCATGCAGGGTGGT-3’ were used. PCR amplification was performed with an initial denaturation at 94°C for 2 min, followed by 34 cycles of denaturation at 94°C for 10 s, annealing at 64°C for 30 s, and extension at 72°C for 30 s, with a final extension at 72°C for 1 min. PCR products were separated by electrophoresis on 1.5% agarose gels and visualized using FAS-BOX3 (NIPPON Genetics). The expected product sizes were 365 bp for the wild-type allele and 375 bp for the *kl*^*sa*18644^ allele (fig. S12).

### Free-swimming behavior recording and analysis

On the morning of behavioral assessment, fish were not fed and were removed from the rearing system. Each fish was transferred together with its rearing tank into a white plastic cage and acclimated for 1 h on the experimental bench used for behavioral recording. After acclimation, the tank was surrounded by opaque partitions to completely block visual stimuli from the experimenter during testing. To improve the tracking accuracy of swimming trajectories, a light source (Tviewer A4-420, Trytech) was placed behind the tank to provide backlighting. Free-swimming behavior was recorded for 10 min at a frame rate of 30 fps using a digital video camera (HDR-CX470, Sony) positioned in front of the tank.

The x-y coordinates of each fish were extracted from the recorded videos using the animal-tracking software UMATracker(*30*). Coordinate data were obtained from all frames of each video (18,000 frames, 600 s), and 14 behavioral features were calculated using Python (version 3.14) and the data-processing libraries pandas (version 2.3.3)(*31*) and NumPy (version 2.3.5)(*32*). To evaluate the vertical spatial distribution of fish within the tank, the tank was divided equally into three areas along the y-axis: Top, Middle, and Bottom. The proportion of time spent in each area and the latency to first entry into the Top area (Latency to Top) were calculated. The number of upward crossings of the y-axis threshold defining the Top area was defined as the number of entries into the Top area. For the calculation of velocity- and distance-related features, coordinate data were smoothed to remove tracking noise using a Savitzky– Golay filter implemented in SciPy (version 1.16.3)(*33*), with a window size of 49 and a polynomial order of 1. Based on the smoothed coordinate data, the total travel distance and travel distances along each axis were calculated. The corresponding mean velocities were obtained by dividing these distances by the observation time. Instantaneous velocity at each time point was also calculated, and its temporal variability was quantified as the variance of velocity. The mean and variance of each coordinate component were further calculated as indices of the spatial activity range of zebrafish.

### Morphological measurements

Morphological measurements were performed on the day following behavioral assessment. Zebrafish were anesthetized with 200 mg/L tricaine (MS-222). After anesthesia, each fish was positioned with its head facing left and tail facing right, and whole-body images were captured from the left lateral side using a digital camera (iPhone, Apple Inc.). After imaging, body weight was measured using a top-loading balance (MS6002S/02, METTLER TOLEDO).

Morphological features were extracted from the images using the image analysis software Fiji(*34*). Linear measurements were performed with reference to previous studies(*35, 36*). The measured traits included standard length (SL), total length (TL), body depth (BD), head length (HL), caudal peduncle length (CPL), caudal peduncle depth (CPD), and caudal fin length (CFL), which was defined as the distance from the base to the distal tip of the caudal fin (fig. S13). Based on these measurements, relative morphological indices, including BD/SL, CPD/CPL, HL/SL, and body weight/TL, were calculated.

### Tissue collection and RNA extraction

After morphological measurements, fish were euthanized by immersion in ice-cold water and were then immediately decapitated and dissected. The intestine was isolated from other visceral tissues, including the liver, pancreas, spleen, and gonads, in 1× PBS. The isolated intestine was transferred to a 1.5 mL tube, mixed with 125 µL of TRI Reagent-LS (Molecular Research Center), and homogenized using a BioMasher II (Nippi). An additional 375 µL of TRI Reagent-LS was then added, and the samples were stored at −80°C until further processing.

RNA extraction was performed according to the standard protocol for TRI Reagent-LS. Briefly, thawed samples were subjected to phase separation by adding chloroform. The aqueous phase was collected, and RNA was precipitated with isopropanol and washed with 75% ethanol. To improve RNA recovery and purity, the resulting RNA solution was further reprecipitated using sodium acetate and ethanol. The dried RNA pellet was dissolved in nuclease-free water. RNA concentration was measured using a Qubit 2.0 Fluorometer (Thermo Fisher Scientific), and RNA samples were adjusted to an appropriate concentration with nuclease-free water. For RNA quality control, representative samples randomly selected from each experimental batch were analyzed using an Agilent 2100 Bioanalyzer (Agilent Technologies). All representative samples had RNA integrity number (RIN) values of 7.0 or higher.

### Library preparation and RNA sequencing

RNA-seq libraries were generated following the Lasy-Seq version 1.1 protocol (https://sites.google.com/view/lasy-seq/). Briefly, extracted RNA was first subjected to reverse transcription using SuperScript IV reverse transcriptase together with indexed reverse transcription primers. The resulting cDNA was purified with an equal volume of AMPure XP beads (Beckman Coulter), and double-stranded cDNA was synthesized using ribonuclease H and DNA polymerase I (Enzymatics). RNase T1 treatment was conducted to digest rRNA. The double-stranded cDNA was then fragmented using 5× WGS Fragmentation Mix (Enzymatics), followed by adapter ligation with 5× WGS Ligase (Enzymatics). Library amplification was carried out using KAPA HiFi HotStart ReadyMix (KAPA Biosystems).

The quality and size distribution of the prepared libraries were examined using an Agilent 2100 Bioanalyzer with a High Sensitivity DNA Kit (Agilent Technologies). Sequencing was performed on a NovaSeq X Plus platform (Illumina), generating 150 bp paired-end reads.

### Data preprocessing and gene quantification

Raw sequencing reads were quality-checked and adapter-trimmed using fastp (version 0.21.0) (*37*). The processed reads were aligned to the zebrafish reference transcriptome of Danio_rerio.GRCz11.101 using BWA-MEM (version 0.7.17) (*38*). Gene-level read counts were then obtained using Salmon (version 0.12.0) (*39*). Gene-level quantification was conducted by aggregating the quantifications of individual transcripts for subsequent analysis.

### Transcriptome and functional enrichment analyses

Transcriptome data processing and enrichment analyses were performed in R (version 4.3.2, https://www.r-project.org/). Differentially expressed genes (DEGs) associated with chronological age were identified using the R package DESeq2 (version 1.42.1)(*40*). Pairwise comparisons were performed among the 4-, 5-, 6-, 7-, and 8-mpf chronological age groups, and genes with an adjusted p-value < 0.05 and |log_2_(Fold Change)| ≥ 1.0 were defined as significant DEGs.

Read count data were further processed using the R package edgeR (version 4.0.16)(*41*). Normalization factors were calculated using the trimmed mean of M-values (TMM) method, and raw counts were transformed into log-transformed counts per million (logCPM). To capture age-associated expression trends, logCPM values were averaged within each chronological age group before clustering. Time-course clustering was performed using fuzzy c-means clustering implemented in the R package Mfuzz (version 2.62.0)(*42*). The fuzzification parameter *m* was estimated using the mestimate function, and the number of clusters was set to 20.

Functional enrichment analyses were performed using the R packages clusterProfiler (version 4.10.1)(*43*) and org.Dr.eg.db (version 3.18.0)(*44*). Genes assigned to each Mfuzz cluster were used as input gene sets, and all genes with detectable expression were used as the background universe. Over-representation analysis (ORA) was performed for Gene Ontology Biological Process (GO BP) terms and KEGG pathways. GO BP enrichment was assessed using zebrafish gene annotations provided by org.Dr.eg.db. KEGG pathway enrichment was assessed using KEGG pathway annotations for Danio rerio with the KEGG organism code “dre”. For KEGG analyses, zebrafish gene symbols were converted to Entrez Gene IDs using org.Dr.eg.db. Enrichment was tested using the hypergeometric test, followed by Benjamini–Hochberg multiple-testing correction. Terms or pathways with an adjusted p-value < 0.05 were considered significant.

To identify molecular pathways associated with non-invasive biological age estimates, gene set enrichment analysis (GSEA) was performed using a ranked gene list based on Spearman’s rank correlation coefficients between gene expression levels, represented as logCPM values, and the InferAging-BM-derived biological age estimate,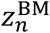. KEGG pathway GSEA was performed using clusterProfiler::gseKEGG with the KEGG organism code “dre” for Danio rerio. Zebrafish gene symbols were converted to Entrez Gene IDs using org.Dr.eg.db before GSEA. Multiple testing correction was performed using the Benjamini–Hochberg method, and gene sets with q-value < 0.10 were considered significant.

### Construction of InferAging-BMT

InferAging-BMT (Behavior, Morphology and Transcriptome) was constructed as a generative model to estimate the latent biological age of each individual by integrating chronological age, sex, behavioral features, morphological features, and transcriptomic features. Chronological age was first standardized to have zero mean and unit variance. Behavioral and morphological features were also z-score standardized. For the transcriptomic modality, Mfuzz(*42*)-derived cluster-level expression features from the intestine were used as sample-level transcriptomic features and were standardized before model fitting.

For individual *n* , the standardized chronological age was denoted by *a*_*n*_ , the binary sex covariate by *s*_*n*_ , the behavioral features by *x*_*n*_, the morphological features by *p*_*n*_ , and the transcriptomic features by *g*_*n*_. The latent biological age, *z*_*n*_, was defined on the standardized chronological-age scale as the sum of chronological age and an individual-specific aging deviation term, *δ*_*n*_:

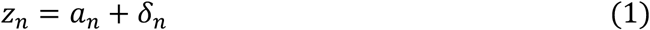

The individual-specific deviation was represented using a non-centered parameterization. An unobserved individual-specific factor, *e*_*n*_, and the deviation term, *δ*_*n*_, were defined as

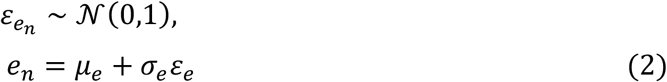

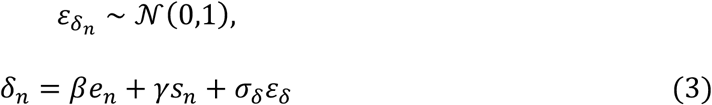

Here, *e*_*n*_ represents an unobserved individual-specific factor contributing to inter-individual variation in aging, *β* represents its effect on the aging deviation, *γ* represents the sex effect, and *σ*_*δ*_ represents residual variation in the deviation term. Although *e*_*n*_ was not directly measured or interpreted in the present study, it was included as a latent individual-specific factor to provide a flexible framework for future extensions incorporating measured environmental covariates and more complex environmental conditions. The priors for these latent components and hyperparameters were specified as

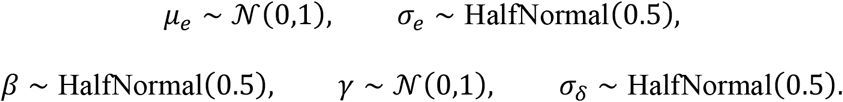

Observed multimodal data were modeled as conditionally dependent on *z*_*n*_. For behavioral feature *j*, morphological feature *k*, and transcriptomic feature *l*, the observation models were defined as

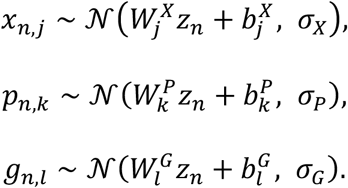

Behavioral and morphological weights were assigned standard normal priors,

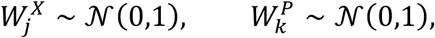

whereas transcriptomic weights were assigned a hierarchical prior,

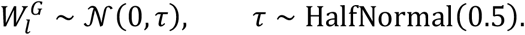

The bias terms were assigned weakly informative priors

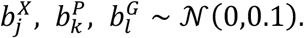

To reduce overfitting under the limited sample size, observation noise was shared across features within each modality. We assigned a gamma distribution as the prior distribution for the precision parameter and defined the corresponding standard deviation as the inverse square root of the precision parameter:

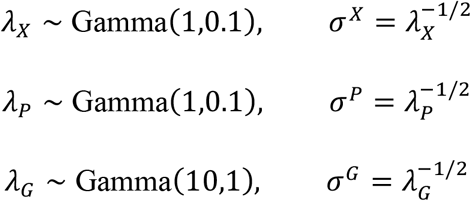

The model was implemented and fitted in Python (version 3.14) using NumPyro (version 0.20.0)(*45*), which uses JAX (version 0.9.0)(*46*) as a backend for accelerated computation. Posterior distributions were estimated using the No-U-Turn Sampler (NUTS), a Markov chain Monte Carlo (MCMC) algorithm. Three independent MCMC chains were run, each with 20,000 warm-up steps and 15,000 sampling steps, yielding a total of 45,000 posterior samples. Convergence was assessed for all parameters using the Gelman–Rubin statistic, with R-hat values < 1.1, calculated using ArviZ (version 0.23.3)(*47*). The final biological age estimate for each individual was defined as the posterior mean, corresponding to the expected a posteriori (EAP) estimate.

### Evaluation of InferAging-BMT using repeated cross-validation

To evaluate the ability of InferAging-BMT to infer biological age in unseen individuals using only non-invasive behavioral and morphological features, repeated 5-fold cross-validation was performed with 100 repetitions. To prevent data leakage due to data splitting, z-score standardization parameters for each feature were calculated using only the training set in each fold and were then applied to the corresponding test set.

In the training phase, the NUTS algorithm described above was applied to the training data using all observed modalities, including behavioral *x*_*n*_, morphological *p*_*n*_, and transcriptomic *g*_*n*_ data. MCMC sampling was performed with 20,000 warm-up steps and 15,000 sampling steps across three chains. The posterior means of the global parameters, including feature weights *W*, bias terms *b*, modality-specific standard deviations *σ*, and the hyperparameters *γ* and *β*, were then calculated and fixed.

In the prediction phase, transcriptomic data, *g*_*n*_, were masked for the test individuals, and only behavioral and morphological data, *x*_*n*_ and *p*_*n*_, were provided to the model. Under this setting, the fixed global parameters estimated from the training data were substituted into the model, and posterior distributions of the local latent variables for the unseen test individuals, including *e*_*n*_ , *δ*_*n*_ , and *z*_*n*_ were newly sampled. This conditional inference step was performed with 20,000 warm-up steps and 20,000 sampling steps across three chains.

For each cross-validation repetition, predictions from folds in which poor MCMC convergence was detected during either the training or prediction phase, defined as 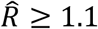, were automatically excluded from the evaluation. The final prediction score for each unseen individual was defined as the average of the posterior means obtained across cross-validation repetitions that satisfied the convergence criterion.

### Construction and evaluation of InferAging-BM

To test whether biological age could be estimated without relying on transcriptomic data, we constructed a fully non-invasive model, InferAging-BM (Behavior and Morphology). InferAging-BM retained the latent-age structure of InferAging-BMT but removed the transcriptomic observation model. Thus, the transcriptomic observation term, *g*_*n*_ , and its associated parameters, *W*^*G*^ , *b*^*G*^ , *σ*^*G*^ , and *τ* , were not included during model fitting. The model was then fitted de novo using all behavioral and morphological features, *X* and *P*, from all individuals as observed data.

For InferAging-BM, parameter estimation by MCMC was performed using the same NUTS settings as those used for InferAging-BMT. The posterior mean of the latent biological age inferred by InferAging-BM for each individual was denoted as 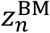 . These estimates were compared with the reference estimates, 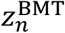 , derived from InferAging-BMT fitted using all modalities, *X, P*, and *G*. The corresponding aging deviation estimates, 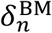 and 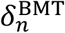, were also compared to evaluate whether inter-individual differences in aging were preserved in the fully non-invasive model.

### One-age-group dropout analysis

To assess whether InferAging-BM estimates were dependent on any single chronological age group, we performed a one-age-group dropout sensitivity analysis. First, InferAging-BM was fitted using all individuals to obtain full-data baseline estimates of biological age 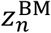 and aging deviation 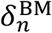. Chronological age groups were assigned according to the nearest representative sampling age corresponding to 4, 5, 6, 7, and 8 mpf. Then, one chronological age group was excluded at a time, and InferAging-BM was refitted using the remaining individuals. For each dropout condition, chronological age, behavioral features, and morphological features were standardized using only the individuals retained in that condition. Posterior means of 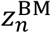 and 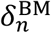 were converted back to the day scale using the corresponding chronological-age scaler.

For each dropout condition, estimates from the dropout model were compared with the full-data baseline estimates for the same retained individuals. Agreement was evaluated using Pearson’s correlation coefficient (*r*), the coefficient of determination (*R*^2^), mean absolute error (MAE), and the sign consistency of 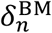, defined as the fraction of individuals for which the sign of 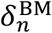 was consistent between the full-data and dropout models.

### Age-imbalance resampling analysis

To evaluate whether InferAging-BM estimates were affected by imbalanced sample sizes across chronological age groups, we performed an age-imbalance resampling analysis. As a baseline, InferAging-BM was first fitted using all individuals to obtain full-data estimates of 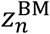 and 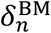. We generated 30 randomly imbalanced subsets by sampling individuals without replacement within each chronological age group while retaining all age groups. Candidate subsets were retained only when they included at least one individual from each age group, contained at least 12 individuals in total, and had a difference of at least two individuals between the largest and smallest age groups. Subsets identical to the full dataset were not allowed.

For each resampled subset, InferAging-BM was refitted de novo using only the individuals included in that subset. Chronological age, behavioral features, and morphological features were standardized within each subset before model fitting. Posterior means of 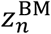 and 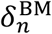 were converted back to the day scale using the chronological-age scaler fitted within each subset. Estimates from each resampled model were compared with the full-data baseline estimates for the same individuals included in that subset. Agreement was evaluated using Pearson’s correlation coefficient (*r*), (*R*^2^), MAE, and the sign consistency of 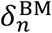. MCMC divergences were recorded for diagnostic purposes.

### Statistical analysis and data visualization

Differences in behavioral and morphological features among chronological age groups were evaluated using the Kruskal–Wallis test implemented in SciPy. When the Kruskal–Wallis test indicated a significant overall difference among age groups (p < 0.05), post hoc pairwise comparisons were performed using Dunn’s test implemented in scikit-posthocs (version 0.12.0)(*48*), with Benjamini–Hochberg false discovery rate (FDR) correction. Pairwise differences with FDR-adjusted q-values < 0.05 were considered significant. The Kruskal– Wallis (p)-values and FDR-adjusted Dunn’s test results are summarized in table S1.

Data splitting, feature scaling, and calculation of model evaluation metrics in cross-validation were implemented using scikit-learn (version 1.7.2)(*49*). For correlation and regression analyses, the normality of residuals was assessed using the Shapiro–Wilk test implemented in SciPy. After confirming that the residuals did not significantly deviate from a normal distribution (p > 0.05), relationships between variables were analyzed using Pearson’s product-moment correlation analysis and linear regression analysis. The predictive accuracy and generalization performance of the biological age estimation models were evaluated by calculating Pearson’s product-moment correlation coefficient (*r*), the coefficient of determination (*R*^2^), and the mean absolute error (MAE).

In Bayesian analyses, posterior means were used as point estimates. Estimation uncertainty was quantified using 95% equal-tailed credible intervals (CIs), defined by the 2.5th and 97.5th percentiles of the posterior distribution. Effects whose 95% CIs did not include zero were considered to have credible non-zero contributions.

To visualize the distribution of high-dimensional data, including behavioral, morphological, and transcriptomic features, Uniform Manifold Approximation and Projection (UMAP) was performed using the Python package umap-learn (version 0.5.9)(*50*). Before dimensionality reduction, all features were standardized using z-score normalization. UMAP was performed with the following parameters: n_neighbors = 10, min_dist = 0.01, and metric = “euclidean”.

## Acknowledgments

We thank Maiko Yokouchi and Yuka Ueda for technical assistance with zebrafish experiments.

## Funding

This work was supported by JSPS KAKENHI Grant Numbers JP21K19265(T.M.), JP22H02821(T.M.), JP23K24083 (T.M.), JP23H02677(Y.B.), the Takeda Science Foundation

and Suntory Foundation for Life Sciences (T.M.), the G-7 Scholarship Foundation; and JST SPRING Grant Number JPMJSP2140 (Y.A.). Moonshot R&D (JPMJMS2024 to H.N.) and CREST (JPMJCR25Q2 to H.N.) from the Japan Science and Technology Agency (JST), Multidisciplinary Frontier Brain and Neuroscience Discoveries (Brain/MINDS 2.0) (JP25wm0625322 to H.N.) from Japan Agency for Medical Research and Development (AMED). AMED Brain/MINDS 2.0 Grant Number JP24wm0625416 (Y.Y.).

## Data and code availability

RNA-sequencing and quantification data generated in this study have been deposited in the Gene Expression Omnibus (GEO) under accession number GSE335259. These data will be made publicly available upon publication.

The custom code used for hierarchical Bayesian modeling and downstream analyses will be made publicly available upon publication.

## Author contributions

Y.A. performed experiments, analyzed data, and developed InferAging frameworks. M.K. performed RNA-seq analyses. H.H. provided the *klotho*-mutant zebrafish line. Y.A., Y.Y., Y.B., H.N., and T.M. conceived the study and acquired funding. Y.A., Y.Y., H.N., and T.M. designed the framework and wrote the manuscript. All authors reviewed and approved the final manuscript.

## Competing interests

The authors declare no competing interests.

## Notes

### Competing Interest Statement

The authors have declared no competing interest.

### Summary of Updates

This revised version includes changes to the title, Figure 1, and related sections to more clearly highlight the central focus and key message of the study.

## Reference

1. D. J. Lowsky, S. J. Olshansky, J. Bhattacharya, D. P. Goldman, Heterogeneity in Healthy Aging. J. Gerontol. A. Biol. Sci. Med. Sci. 69, 640–649 (2014).

2. A. Mitnitski, S. E. Howlett, K. Rockwood, Heterogeneity of Human Aging and Its Assessment.J. Gerontol. A. Biol. Sci. Med. Sci., glw089 (2016).

3. M. E. Levine, Modeling the Rate of Senescence: Can Estimated Biological Age Predict Mortality More Accurately Than Chronological Age? J. Gerontol. A. Biol. Sci. Med. Sci. 68, 667–674 (2013).

4. D. W. Belsky, A. Caspi, R. Houts, H. J. Cohen, D. L. Corcoran, A. Danese, H. Harrington, S. Israel, M. E. Levine, J. D. Schaefer, K. Sugden, B. Williams, A. I. Yashin, R. Poulton, T. E. Moffitt, Quantification of biological aging in young adults. Proc. Natl. Acad. Sci. 112 (2015).

5. J. C. Whitehead, B. A. Hildebrand, M. Sun, M. R. Rockwood, R. A. Rose, K. Rockwood, S. E. Howlett, A Clinical Frailty Index in Aging Mice: Comparisons With Frailty Index Data in Humans. J. Gerontol. Ser. A 69, 621–632 (2014).

6. S. Bocklandt, W. Lin, M. E. Sehl, F. J. Sánchez, J. S. Sinsheimer, S. Horvath, E. Vilain, Epigenetic Predictor of Age. PLoS ONE 6, e14821 (2011).

7. S. Horvath, DNA methylation age of human tissues and cell types. Genome Biol. 14, 3156 (2013).

8. J. Rutledge, H. Oh, T. Wyss-Coray, Measuring biological age using omics data. Nat. Rev. Genet. 23, 715–727 (2022).

9. C. Huang, C. Xiong, K. Kornfeld, Measurements of age-related changes of physiological processes that predict lifespan of Caenorhabditis elegans. Proc. Natl. Acad. Sci. 101, 8084– 8089 (2004).

10. K. G. Iliadi, G. L. Boulianne, Age-related behavioral changes in Drosophila. Ann. N. Y. Acad. Sci. 1197, 9–18 (2010).

11. H. Shoji, K. Takao, S. Hattori, T. Miyakawa, Age-related changes in behavior in C57BL/6J mice from young adulthood to middle age. Mol. Brain 9, 11 (2016).

12. A. Turturro, W. W. Witt, S. Lewis, B. S. Hass, R. D. Lipman, R. W. Hart, Growth Curves and Survival Characteristics of the Animals Used in the Biomarkers of Aging Program. J. Gerontol.A. Biol. Sci. Med. Sci. 54, B492–B501 (1999).

13. C. Pettan-Brewer, PiperM. M. Treuting, Practical pathology of aging mice. Pathobiol. Aging Age-Relat. Dis. 1, 7202 (2011).

14. A. Fahlström, Q. Yu, B. Ulfhake, Behavioral changes in aging female C57BL/6 mice. Neurobiol. Aging 32, 1868–1880 (2011).

15. M. J. H. Gilbert, T. C. Zerulla, K. B. Tierney, Zebrafish (Danio rerio) as a model for the study of aging and exercise: Physical ability and trainability decrease with age. Exp. Gerontol. 50, 106–113 (2014).

16. L. Yu, V. Tucci, S. Kishi, I. V. Zhdanova, Cognitive Aging in Zebrafish. PLoS ONE 1, e14 (2006).

17. G. S. Gerhard, E. J. Kauffman, X. Wang, R. Stewart, J. L. Moore, C. J. Kasales, E. Demidenko,K. C. Cheng, Life spans and senescent phenotypes in two strains of Zebrafish (Danio rerio).Exp. Gerontol. 37, 1055–1068 (2002).

18. G. S. Gerhard, K. C. Cheng, A call to fins! Zebrafish as a gerontological model. Aging Cell 1, 104–111 (2002).

19. J. Hudock, J. W. Kenney, Aging in zebrafish is associated with reduced locomotor activity and strain dependent changes in bottom dwelling and thigmotaxis. PLOS ONE 19, e0300227 (2024).

20. C. N. Bedbrook, R. D. Nath, L. Zhang, S. W. Linderman, A. Brunet, K. Deisseroth, Lifelong behavioral screen reveals an architecture of vertebrate aging. Science 391, eaea9795 (2026).

21. Q. Zhang, C. L. Vallerga, R. M. Walker, T. Lin, A. K. Henders, G. W. Montgomery, J. He, D. Fan, J. Fowdar, M. Kennedy, T. Pitcher, J. Pearson, G. Halliday, J. B. Kwok, I. Hickie, S. Lewis,T. Anderson, P. A. Silburn, G. D. Mellick, S. E. Harris, P. Redmond, A. D. Murray, D. J. Porteous, C. S. Haley, K. L. Evans, A. M. McIntosh, J. Yang, J. Gratten, R. E. Marioni, N. R. Wray, I. J. Deary, A. F. McRae, P. M. Visscher, Improved precision of epigenetic clock estimates across tissues and its implication for biological ageing. Genome Med. 11, 54 (2019).

22. C. G. Bell, R. Lowe, P. D. Adams, A. A. Baccarelli, S. Beck, J. T. Bell, B. C. Christensen, V. N. Gladyshev, B. T. Heijmans, S. Horvath, T. Ideker, J.-P. J. Issa, K. T. Kelsey, R. E. Marioni, W. Reik, C. L. Relton, L. C. Schalkwyk, A. E. Teschendorff, W. Wagner, K. Zhang, V. K. Rakyan, DNA methylation aging clocks: challenges and recommendations. Genome Biol. 20, 249 (2019).

23. E. Bernabeu, D. L. McCartney, D. A. Gadd, R. F. Hillary, A. T. Lu, L. Murphy, N. Wrobel, A. Campbell, S. E. Harris, D. Liewald, C. Hayward, C. Sudlow, S. R. Cox, K. L. Evans, S. Horvath,A. M. McIntosh, M. R. Robinson, C. A. Vallejos, R. E. Marioni, Refining epigenetic prediction of chronological and biological age. Genome Med. 15, 12 (2023).

24. Y. Ogura, R. Kaneko, K. Ujibe, Y. Wakamatsu, H. Hirata, Loss of αklotho causes reduced motor ability and short lifespan in zebrafish. Sci. Rep. 11, 15090 (2021).

25. C. López-Otín, M. A. Blasco, L. Partridge, M. Serrano, G. Kroemer, Hallmarks of aging: An expanding universe. Cell 186, 243–278 (2023).

26. D. Kriukov, E. Efimov, M. S. Gelfand, A. Moskalev, E. E. Khrameeva, Do we actually need aging clocks? Npj Aging 12, 1–9 (2026).

27. L. P. Fried, C. M. Tangen, J. Walston, A. B. Newman, C. Hirsch, J. Gottdiener, T. Seeman, R. Tracy, W. J. Kop, G. Burke, M. A. McBurnie, Frailty in Older Adults: Evidence for a Phenotype.J. Gerontol. A. Biol. Sci. Med. Sci. 56, M146–M157 (2001).

28. D. Yılmaz, A. Singh, E. Wehrle, G. A. Kuhn, N. Mathavan, R. Müller, Unveiling frailty: comprehensive and sex-specific characterization in prematurely aging PolgA mice. Front. Aging 5, 1365716 (2024).

29. C. Takahashi, E. Okabe, M. Nono, S. Kishimoto, H. Matsui, T. Ishitani, T. Yamamoto, M. Uno,E. Nishida, Single housing of juveniles accelerates early-stage growth but extends adult lifespan in African turquoise killifish. Aging 16, 12443–12472 (2024).

30. O. Yamanaka, R. Takeuchi, UMATracker: an intuitive image-based tracking platform. J. Exp. Biol., jeb.182469 (2018).

31. T. pandas development team, pandas-dev/pandas: Pandas, version latest, Zenodo (2020); 10.5281/zenodo.3509134.

32. C. R. Harris, K. J. Millman, S. J. van der Walt, R. Gommers, P. Virtanen, D. Cournapeau, E. Wieser, J. Taylor, S. Berg, N. J. Smith, R. Kern, M. Picus, S. Hoyer, M. H. van Kerkwijk, M. Brett, A. Haldane, J. F. del Río, M. Wiebe, P. Peterson, P. Gérard-Marchant, K. Sheppard, T. Reddy, W. Weckesser, H. Abbasi, C. Gohlke, T. E. Oliphant, Array programming with NumPy. Nature 585, 357–362 (2020).

33. P. Virtanen, R. Gommers, T. E. Oliphant, M. Haberland, T. Reddy, D. Cournapeau, E. Burovski, P. Peterson, W. Weckesser, J. Bright, S. J. van der Walt, M. Brett, J. Wilson, K. J. Millman, N. Mayorov, A. R. J. Nelson, E. Jones, R. Kern, E. Larson,İ. Polat, Y. Feng, E. W. Moore, J. VanderPlas, D. Laxalde, J. Perktold, R. Cimrman, I. Henriksen, E. A. Quintero, C. R. Harris, A. M. Archibald, A. H. Ribeiro, F. Pedregosa, P. van Mulbregt, SciPy1.0 Contributors, SciPy 1.0: Fundamental Algorithms for Scientific Computing in Python. Nat. Methods 17, 261–272 (2020).

34. J. Schindelin, I. Arganda-Carreras, E. Frise, V. Kaynig, M. Longair, T. Pietzsch, S. Preibisch,C. Rueden, S. Saalfeld, B. Schmid, J.-Y. Tinevez, D. J. White, V. Hartenstein, K. Eliceiri, P. Tomancak, A. Cardona, Fiji: an open-source platform for biological-image analysis. Nat. Methods 9, 676–682 (2012).

35. A. Oikonomou, F. Leprieur, I. D. Leonardos, Ecomorphological diversity of freshwater fishes as a tool for conservation priority setting: a case study from a Balkan hotspot. Environ. Biol. Fishes 101, 1121–1136 (2018).

36. X. Cao, J. Zhao, C. Li, S. Zhu, Y. Hao, Y. Cheng, H. Wu, Morphological and skeletal comparison and ecological adaptability of Mandarin fish Siniperca chuatsi and big-eye Mandarin fish Siniperca kneri. Aquac. Fish. 6, 455–464 (2021).

37. S. Chen, Y. Zhou, Y. Chen, J. Gu, fastp: an ultra-fast all-in-one FASTQ preprocessor.Bioinformatics 34, i884–i890 (2018).

38. H. Li, R. Durbin, Fast and accurate short read alignment with Burrows–Wheeler transform.Bioinformatics 25, 1754–1760 (2009).

39. R. Patro, G. Duggal, M. I. Love, R. A. Irizarry, C. Kingsford, Salmon provides fast and bias-aware quantification of transcript expression. Nat. Methods 14, 417–419 (2017).

40. M. I. Love, W. Huber, S. Anders, Moderated estimation of fold change and dispersion for RNA-seq data with DESeq2. Genome Biol. 15, 550 (2014).

41. Y. Chen, L. Chen, A. T. L. Lun, P. L. Baldoni, G. K. Smyth, edgeR v4: powerful differential analysis of sequencing data with expanded functionality and improved support for small counts and larger datasets. Nucleic Acids Res. 53, gkaf018 (2025).

42. M. E. Futschik, B. Carlisle, NOISE-ROBUST SOFT CLUSTERING OF GENE EXPRESSION TIME-COURSE DATA. J. Bioinform. Comput. Biol. 03, 965–988 (2005).

43. T. Wu, E. Hu, S. Xu, M. Chen, P. Guo, Z. Dai, T. Feng, L. Zhou, W. Tang, L. Zhan, X. Fu, S. Liu, X. Bo, G. Yu, clusterProfiler 4.0: A universal enrichment tool for interpreting omics data. The Innovation 2, 100141 (2021).

44. M. Carlson, org.Dr.eg.db: Genome wide annotation for Zebrafish (2023). https://bioconductor.org/packages/org.Dr.eg.db/.

45. D. Phan, N. Pradhan, M. Jankowiak, Composable Effects for Flexible and Accelerated Probabilistic Programming in NumPyro. ArXiv Prepr. ArXiv191211554 (2019).

46. J. Bradbury, R. Frostig, P. Hawkins, M. J. Johnson, Y. Katariya, C. Leary, D. Maclaurin, G. Necula, A. Paszke, J. VanderPlas, S. Wanderman-Milne, Q. Zhang, JAX: composable transformations of Python+NumPy programs, version 0.3.13 (2018); http://github.com/jax-ml/jax.

47. R. Kumar, C. Carroll, A. Hartikainen, O. Martin, ArviZ a unified library for exploratory analysis of Bayesian models in Python. J. Open Source Softw. 4, 1143 (2019).

48. M. Terpilowski, scikit-posthocs: Pairwise multiple comparison tests in Python. J. Open Source Softw. 4, 1169 (2019).

49. F. Pedregosa, G. Varoquaux, A. Gramfort, V. Michel, B. Thirion, O. Grisel, M. Blondel, P. Prettenhofer, R. Weiss, V. Dubourg, J. Vanderplas, A. Passos, D. Cournapeau, M. Brucher, M. Perrot, E. Duchesnay, Scikit-learn: Machine Learning in Python. J. Mach. Learn. Res. 12, 2825–2830 (2011).

50. L. McInnes, J. Healy, N. Saul, L. Grossberger, UMAP: Uniform Manifold Approximation and Projection. J. Open Source Softw. 3, 861 (2018).

