## Supplementary material for "InferAging: A non-invasive aging clock for quantifying individual differences in aging": SI

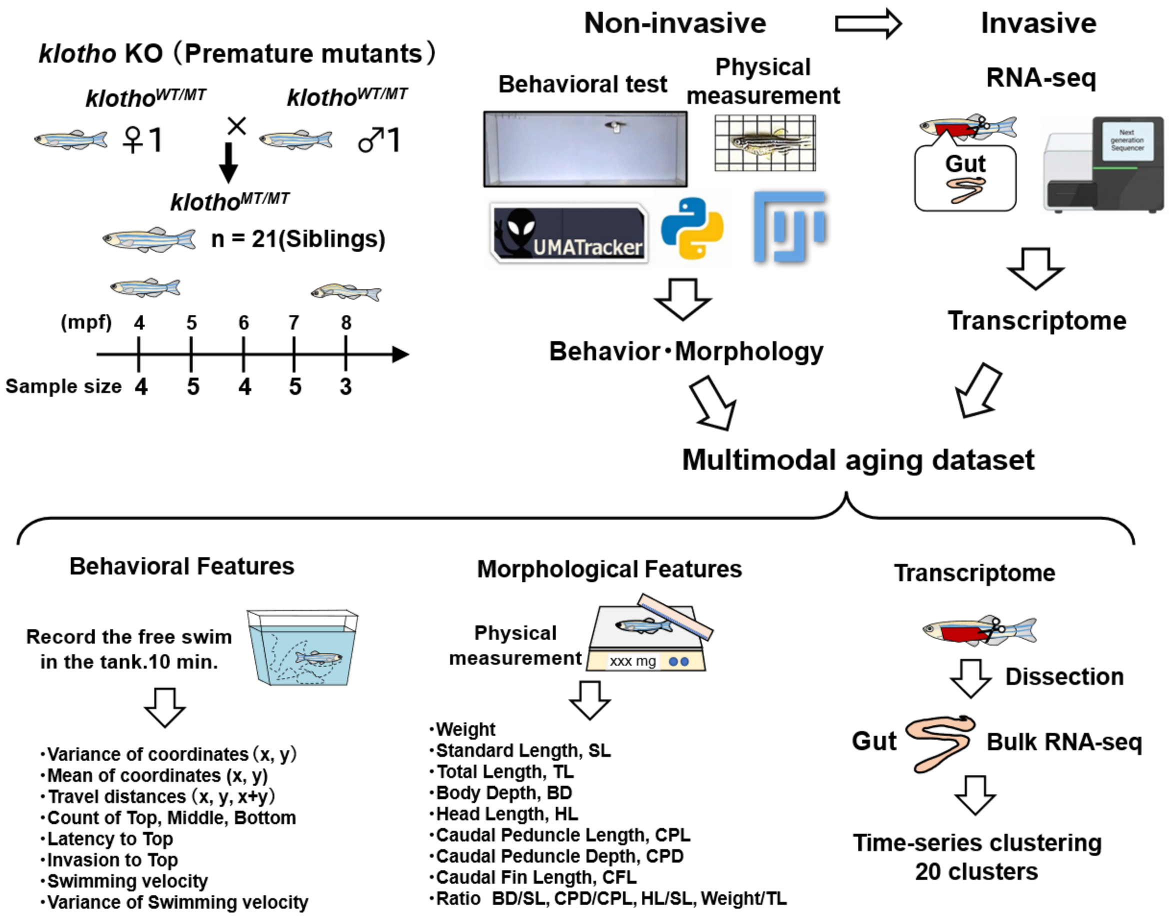


**Fig. S1. Establishment of a multimodal dataset integrating behavioral, morphological, and transcriptomic data.**
Schematic overview of the experimental design and data acquisition. A multimodal dataset consisting of non-invasive behavioral and morphological data, together with intestinal transcriptomic data, was obtained from individual-matched $kl^{-/-}$ zebrafish across five age groups: 4, 5, 6, 7, and 8 months post-fertilization (mpf) (n = 21).

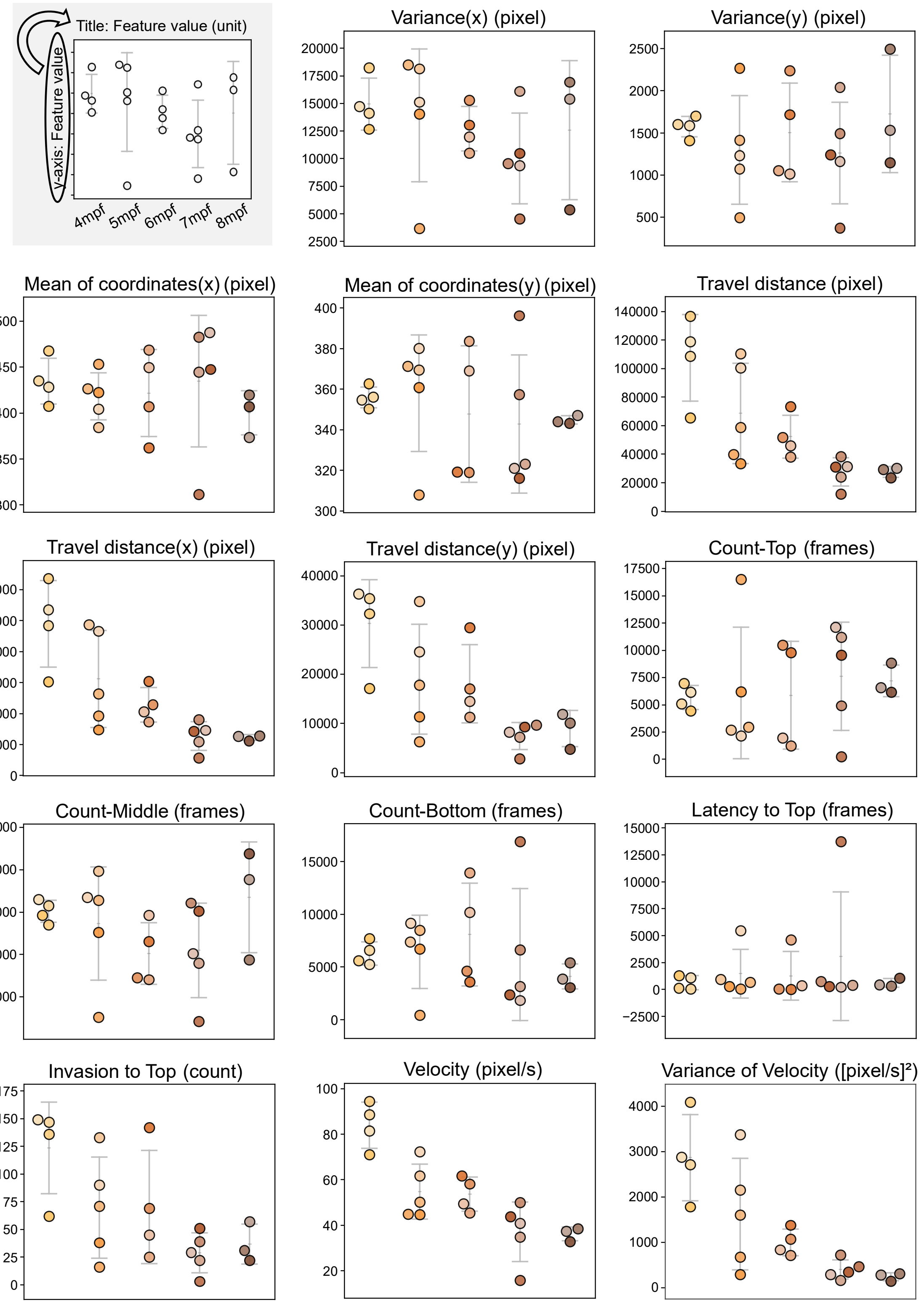


**Fig. S2. Age-associated changes in free-swimming behavioral features in $kl^{-/-}$ zebrafish.** Swarm plots showing the distribution of behavioral indices extracted from free-swimming trajectories of $kl^{-/-}$ zebrafish aged 4–8 mpf (n = 21). The upper-left guide panel schematically illustrates the common layout of each feature panel. Each dot represents an individual fish. Gray horizontal markers and vertical error bars indicate the mean ± s.d. within each age group. The x-axis indicates chronological age in months post-fertilization (mpf). Feature names and corresponding measurement units are shown above each panel.

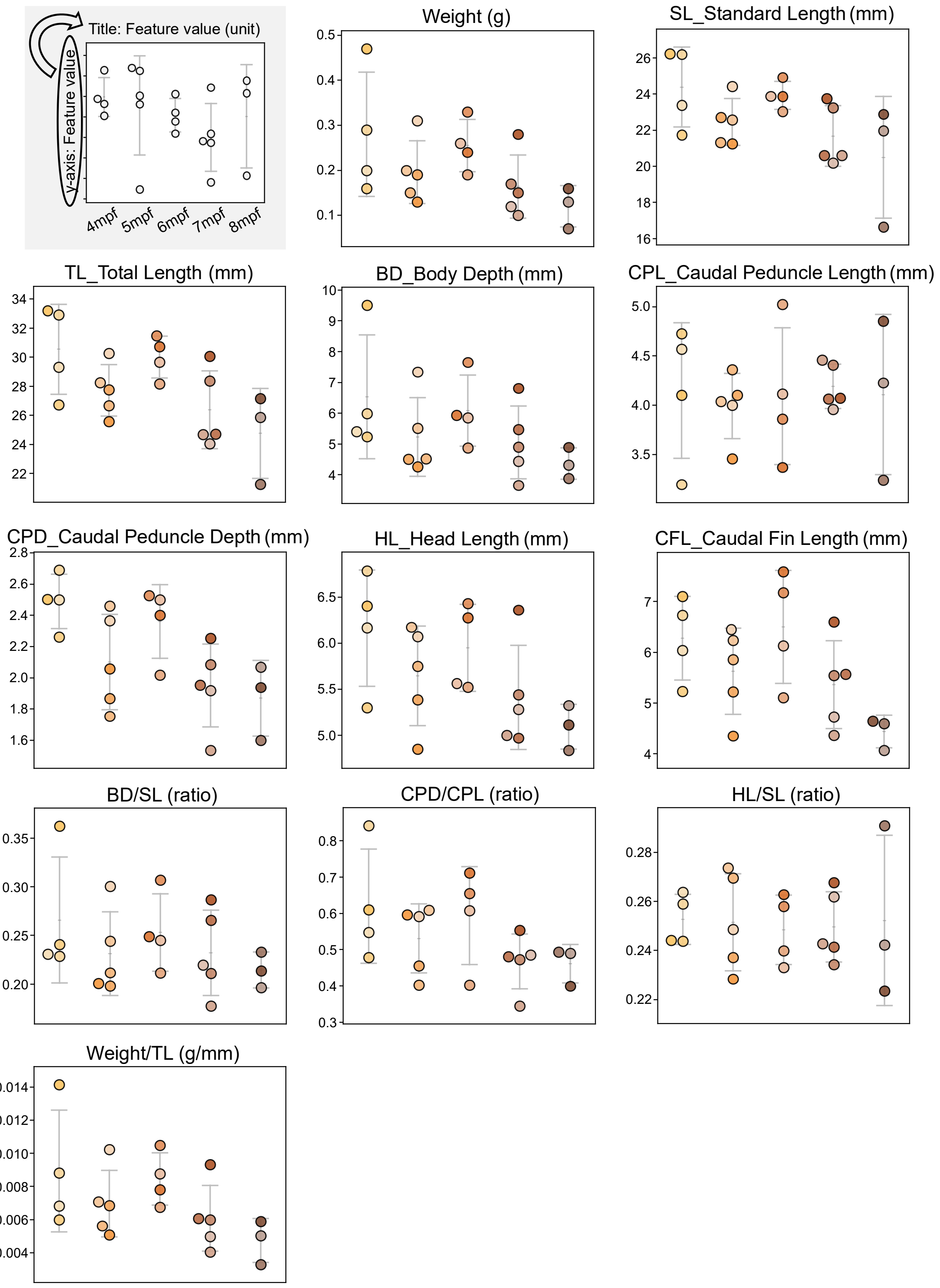


**Fig. S3. Age-associated changes in morphological indices in $kl^{-/-}$ zebrafish.**
Swarm plots showing the distribution of morphological indices obtained from $kl^{-/-}$ zebrafish aged 4–8 mpf (n = 21). The upper-left guide panel schematically illustrates the common layout of each feature panel. Each dot represents an individual fish. Gray horizontal markers and vertical error bars indicate the mean ± s.d. within each age group. The x-axis indicates chronological age in months post-fertilization (mpf). Feature names and corresponding measurement units are shown above each panel.

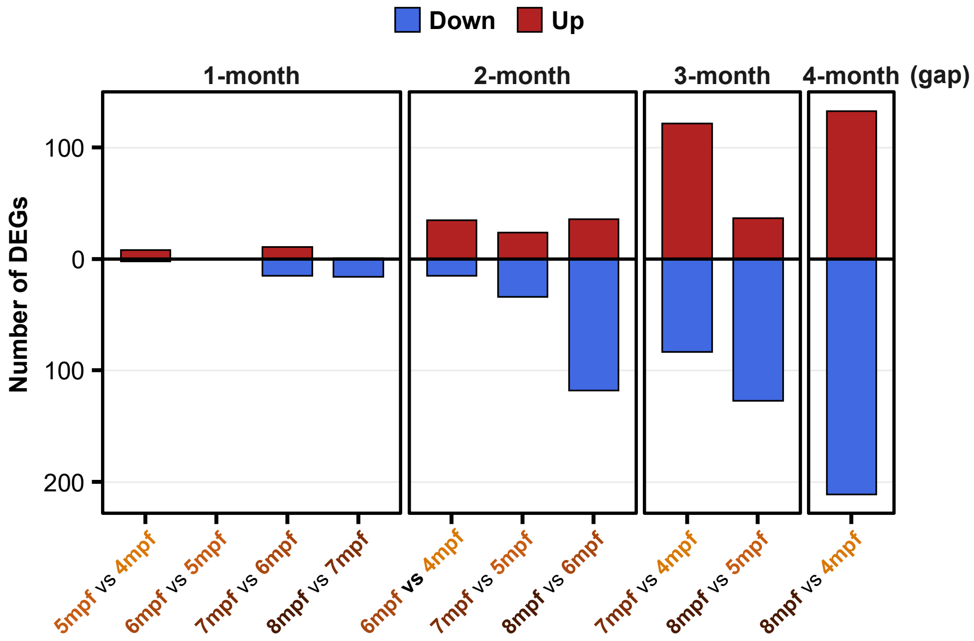


**Fig. S4. Number of differentially expressed genes between age groups in the intestine of $kl^{-/-}$ zebrafish.**

Bar plot summarizing the number of differentially expressed genes (DEGs) between all pairs of age groups based on RNA-seq data obtained from the intestine of $kl^{-/-}$ zebrafish aged 4–8 mpf. The x-axis indicates the compared age-group pairs, and the y-axis indicates the number of detected DEGs. Red bars represent significantly upregulated genes (Up), and blue bars represent significantly downregulated genes (Down). DEGs were identified using thresholds of adjusted p-value < 0.05 and $|\log_2(\text{fold change})| \geq 1.0$.

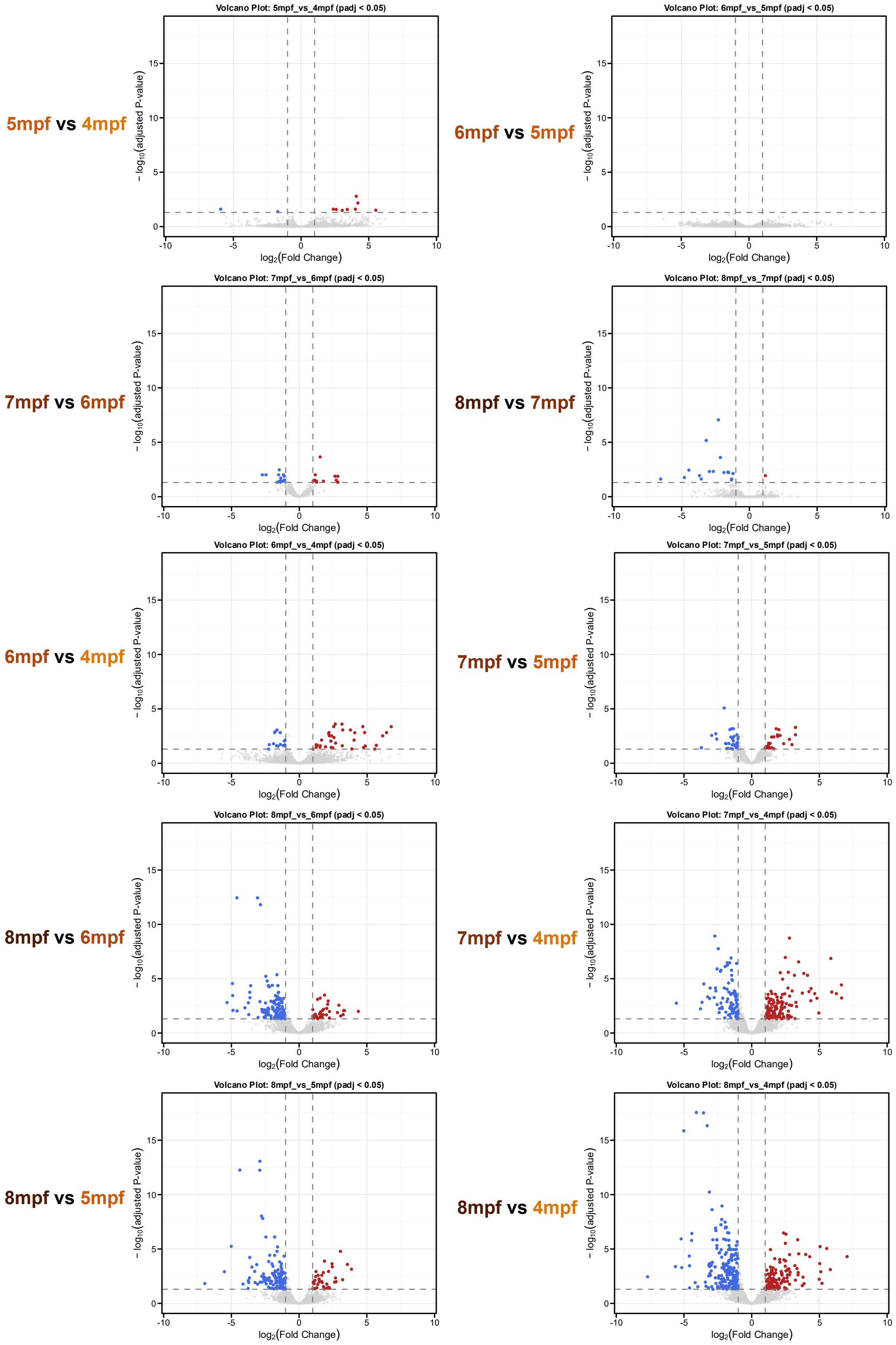

5mpf vs 4mpf
Volcano Plot: 5mpf_vs_4mpf (padj < 0.05)
6mpf vs 5mpf
Volcano Plot: 6mpf_vs_5mpf (padj < 0.05)
7mpf vs 6mpf
Volcano Plot: 7mpf_vs_6mpf (padj < 0.05)
8mpf vs 7mpf
Volcano Plot: 8mpf_vs_7mpf (padj < 0.05)
6mpf vs 4mpf
Volcano Plot: 6mpf_vs_4mpf (padj < 0.05)
7mpf vs 5mpf
Volcano Plot: 7mpf_vs_5mpf (padj < 0.05)
8mpf vs 6mpf
Volcano Plot: 8mpf_vs_6mpf (padj < 0.05)
7mpf vs 4mpf
Volcano Plot: 7mpf_vs_4mpf (padj < 0.05)
8mpf vs 5mpf
Volcano Plot: 8mpf_vs_5mpf (padj < 0.05)
8mpf vs 4mpf
Volcano Plot: 8mpf_vs_4mpf (padj < 0.05)
$-\log_{10}$(adjusted P-value)
$\log_2$(Fold Change)

**Fig. S5. Volcano plots of gene expression changes between age-group pairs.**

Volcano plots showing changes in individual gene expression across all pairwise comparisons among the 4–8 mpf age groups (10 comparisons in total). The x-axis represents $\log_2$(fold change), indicating the magnitude of differential expression, and the y-axis represents $-\log_{10}$(adjusted p-value), indicating statistical significance. Each dot represents a single gene. Red dots indicate significantly upregulated genes (Up), blue dots indicate significantly downregulated genes (Down), and gray dots indicate genes without significant changes. Gray dashed vertical and horizontal lines indicate the DEG detection thresholds: adjusted p-value = 0.05 and $|\log_2(f\text{old change})| \geq 1.0$.

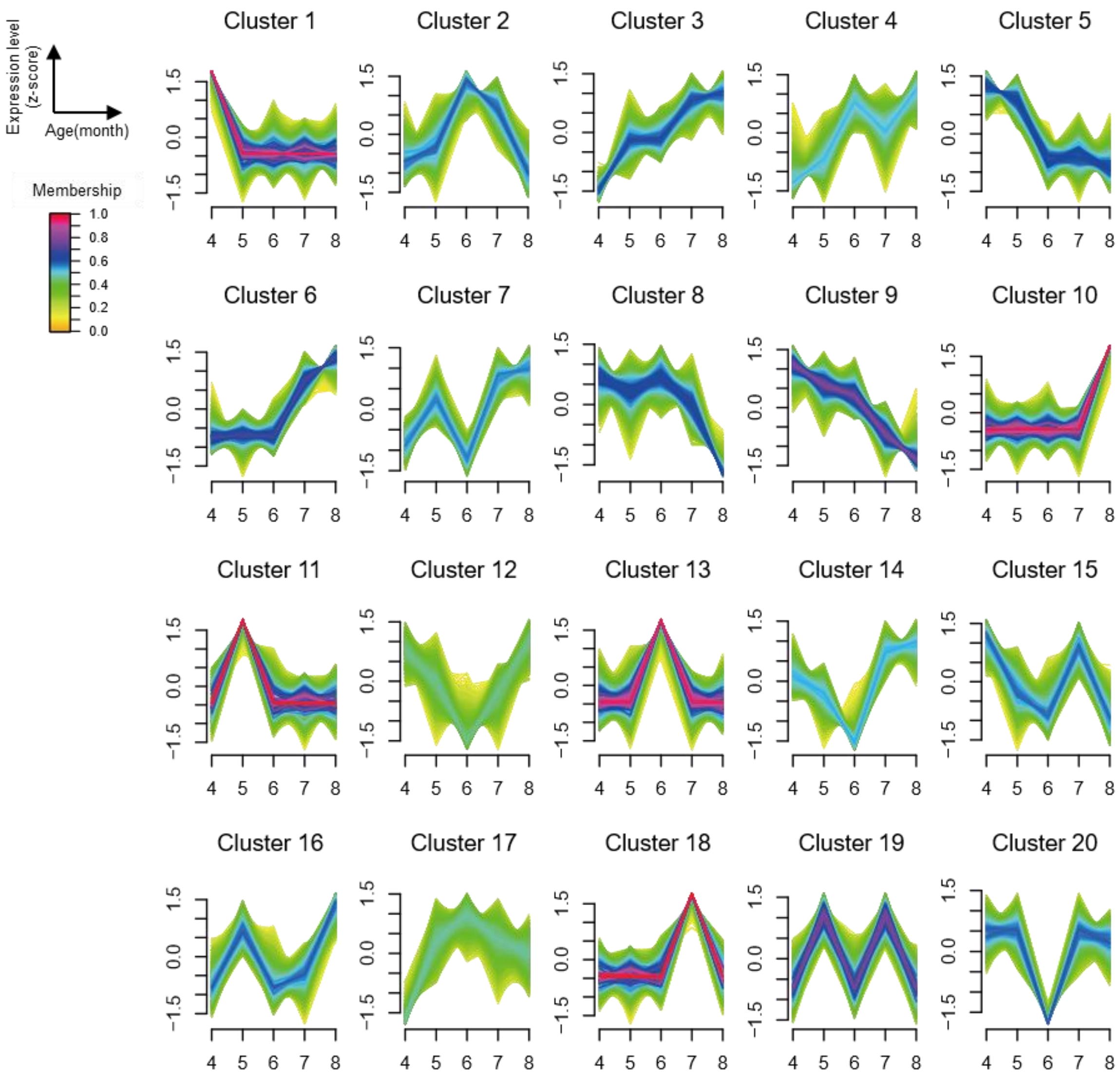


**Fig. S6. Time-series clustering of age-associated gene expression in the intestine of $kl^{-/-}$ zebrafish.**

Results of Mfuzz c-means clustering analysis of transcriptomic data obtained from the intestine of $kl^{-/-}$ zebrafish aged 4–8 mpf. The x-axis indicates age in months post-fertilization (mpf), and the y-axis indicates standardized (z-scored) gene expression levels. By integrating data from all individuals, all genes were classified into 20 co-expression clusters (Clusters 1–20) based on age-associated temporal expression dynamics.

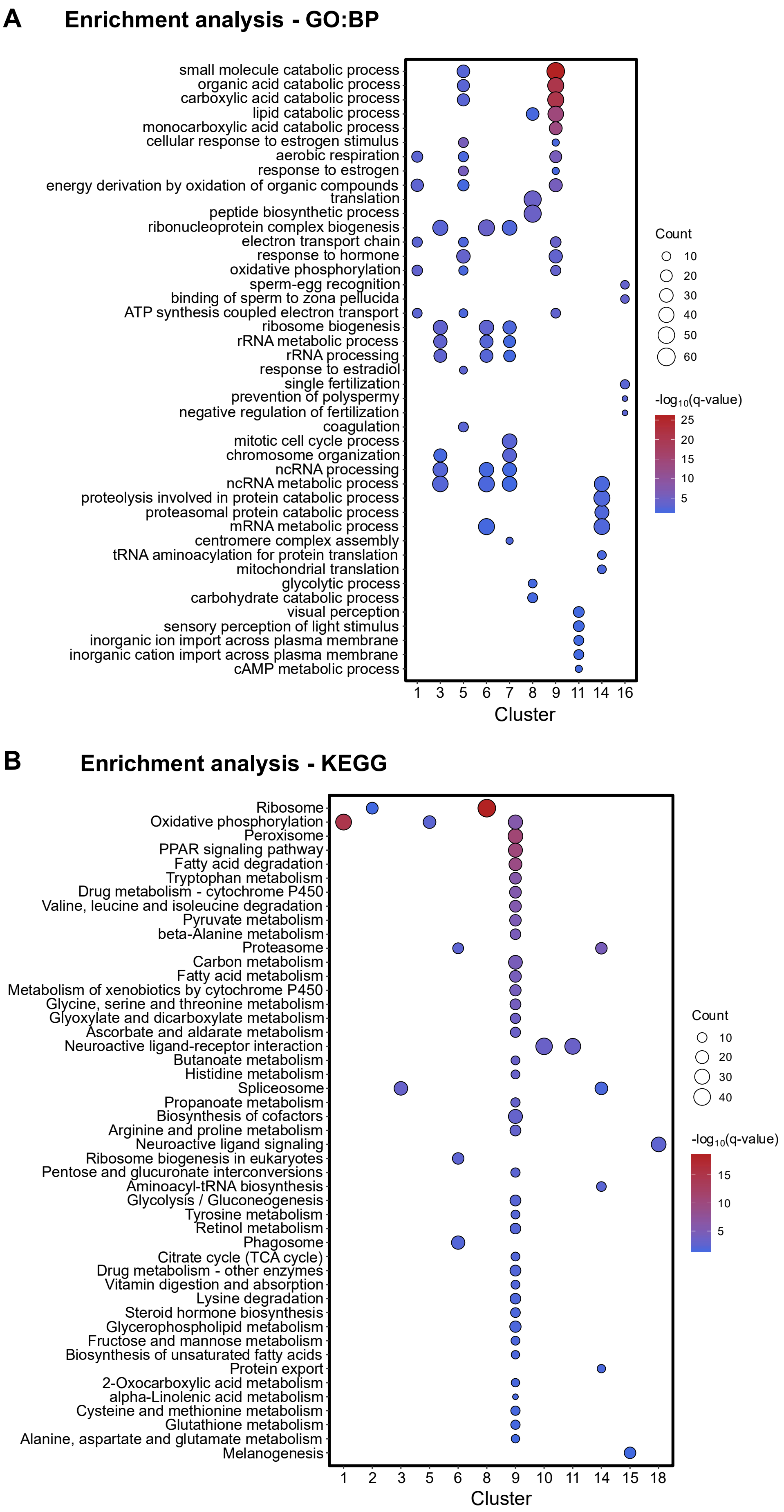
A
Enrichment analysis - GO:BP
small molecule catabolic process
organic acid catabolic process
carboxylic acid catabolic process
lipid catabolic process
monocarboxylic acid catabolic process
cellular response to estrogen stimulus
aerobic respiration
response to estrogen
energy derivation by oxidation of organic compounds
translation
peptide biosynthetic process
ribonucleoprotein complex biogenesis
electron transport chain
response to hormone
oxidative phosphorylation
sperm-egg recognition
binding of sperm to zona pellucida
ATP synthesis coupled electron transport
ribosome biogenesis
rRNA metabolic process
rRNA processing
response to estradiol
single fertilization
prevention of polyspermy
negative regulation of fertilization
coagulation
mitotic cell cycle process
chromosome organization
ncRNA processing
ncRNA metabolic process
proteolysis involved in protein catabolic process
proteasomal protein catabolic process
mRNA metabolic process
centromere complex assembly
tRNA aminoacylation for protein translation
mitochondrial translation
glycolytic process
carbohydrate catabolic process
visual perception
sensory perception of light stimulus
inorganic ion import across plasma membrane
inorganic cation import across plasma membrane
cAMP metabolic process
1 3 5 6 7 8 9 11 14 16
Cluster
Count
10
20
30
40
50
60
-log10(q-value)
25
20
15
10
5
B
Enrichment analysis - KEGG
Ribosome
Oxidative phosphorylation
Peroxisome
PPAR signaling pathway
Fatty acid degradation
Tryptophan metabolism
Drug metabolism - cytochrome P450
Valine, leucine and isoleucine degradation
Pyruvate metabolism
beta-Alanine metabolism
Proteasome
Carbon metabolism
Fatty acid metabolism
Metabolism of xenobiotics by cytochrome P450
Glycine, serine and threonine metabolism
Glyoxylate and dicarboxylate metabolism
Ascorbate and aldarate metabolism
Neuroactive ligand-receptor interaction
Butanoate metabolism
Histidine metabolism
Spliceosome
Propanoate metabolism
Biosynthesis of cofactors
Arginine and proline metabolism
Neuroactive ligand signaling
Ribosome biogenesis in eukaryotes
Pentose and glucuronate interconversions
Aminoacyl-tRNA biosynthesis
Glycolysis / Gluconeogenesis
Tyrosine metabolism
Retinol metabolism
Phagosome
Citrate cycle (TCA cycle)
Drug metabolism - other enzymes
Vitamin digestion and absorption
Lysine degradation
Steroid hormone biosynthesis
Glycerophospholipid metabolism
Fructose and mannose metabolism
Biosynthesis of unsaturated fatty acids
Protein export
2-Oxocarboxylic acid metabolism
alpha-Linolenic acid metabolism
Cysteine and methionine metabolism
Glutathione metabolism
Alanine, aspartate and glutamate metabolism
Melanogenesis
1 2 3 5 6 8 9 10 11 14 15 18
Cluster
Count
10
20
30
40
-log10(q-value)
15
10
5

**Fig. S7. Enrichment analysis of expression clusters obtained by time-series clustering.**
Dot plots showing the results of over-representation analysis (ORA) for expression clusters identified by Mfuzz c-means clustering. Panels (A) and (B) show the results based on Gene Ontology (GO) Biological Process and the KEGG pathway database, respectively. The y-axis indicates significantly enriched GO terms or pathway names. Dot size represents the number of genes included in the corresponding term or pathway, and the color of each dot indicates $-\log_{10}$(q-value).

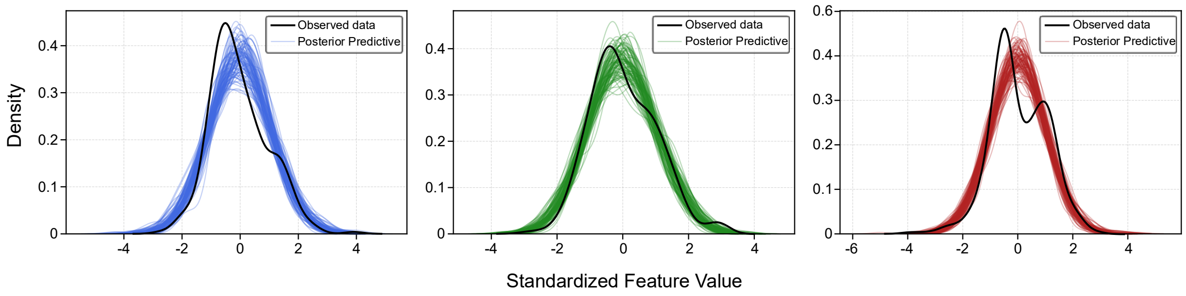


**Fig. S8. Posterior predictive check of InferAging-BMT.**
Comparison between the observed data and posterior predictive distributions of InferAging-BMT, for which parameters were estimated using integrated data from all modalities. Each panel represents one data modality: behavior, morphology, or transcriptome. The x-axis indicates standardized (z-scored) values of each feature, and the y-axis indicates probability density. The thick black solid line represents the distribution of the actual observed data. Colored semi-transparent lines, with blue for behavior, green for morphology, and red for transcriptome, represent distributions of random samples drawn from the posterior predictive distribution of the estimated model.

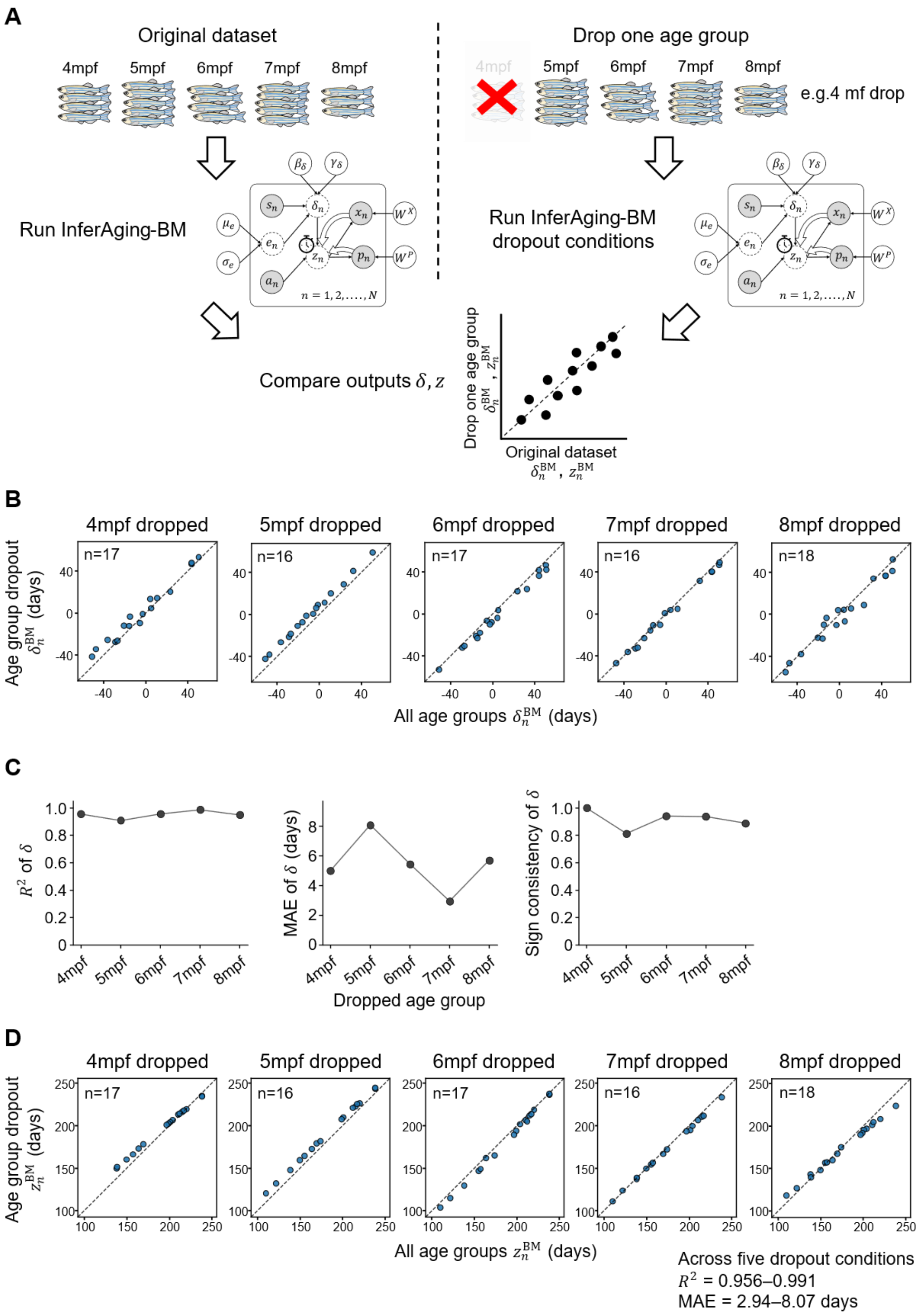


**Fig. S9. One-age-group-dropout sensitivity analysis of InferAging-BM estimates.**
(**A**) Schematic overview of the one-age-group-dropout sensitivity analysis. InferAging-BM was refitted after removing one chronological age group at a time, and estimates for the remaining shared individuals were compared with full-data InferAging-BM estimates. (**B**) Comparison of

aging deviation $\delta_n^{\mathrm{BM}}$ estimates between full-data InferAging-BM and age-group-dropout InferAging-BM across dropout conditions. Each panel represents one removed chronological age group. The dashed line indicates the identity line. (**C**) Summary metrics for the robustness of $\delta_n^{\mathrm{BM}}$ estimates across dropout conditions, including $R^2$, mean absolute error (MAE), and sign consistency. Sign consistency was defined as the proportion of individuals whose $\delta_n^{\mathrm{BM}}$ sign was preserved between full-data and one-age-group-dropout InferAging-BM estimates. (**D**) Comparison of biological age $z_n^{\mathrm{BM}}$ estimates between full-data InferAging-BM and one-age-group-dropout InferAging-BM across dropout conditions. Each panel represents one removed chronological age group. Across the five dropout conditions, $R^2$ for $z_n^{\mathrm{BM}}$ ranged from 0.956 to 0.991, and MAE ranged from 2.94 to 8.07 days.

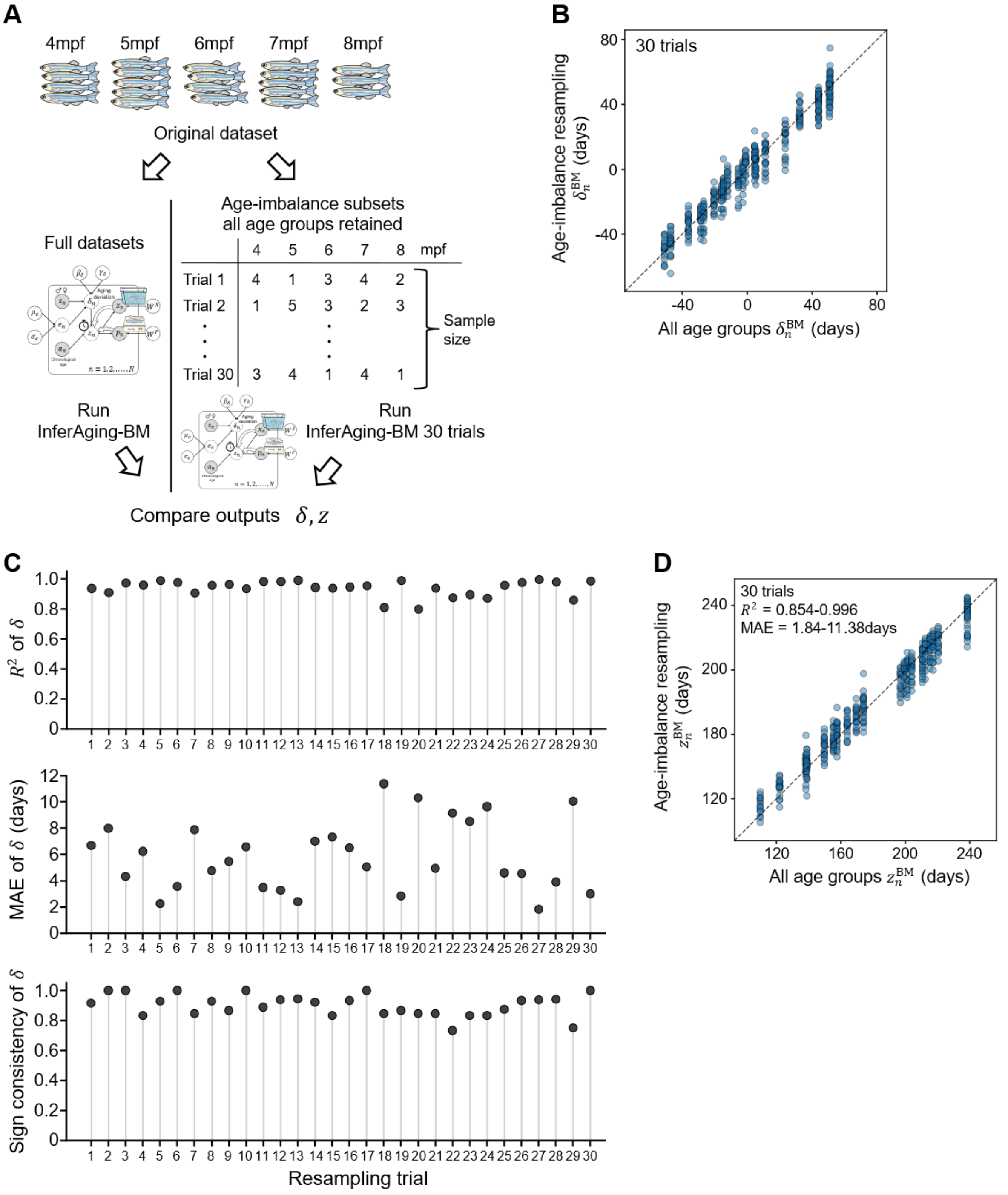


| | 4 | 5 | 6 | 7 | 8 | mpf |
|---|---|---|---|---|---|---|
| Trial 1 | 4 | 1 | 3 | 4 | 2 | |
| Trial 2 | 1 | 5 | 3 | 2 | 3 | |
| ⋮ | | | ⋮ | | | |
| Trial 30 | 3 | 4 | 1 | 4 | 1 | |

**Fig. S10. Age-imbalance resampling analysis of InferAging-BM estimates.**
(**A**) Schematic overview of the age-imbalance resampling analysis. Randomly imbalanced subsets were generated while retaining all chronological age groups, and InferAging-BM was refitted for each subset. Estimates for individuals included in each subset were compared with full-data InferAging-BM estimates. (**B**) Pooled comparison of aging deviation $\delta_n^{\mathrm{BM}}$ estimates between full-data InferAging-BM and age-imbalance-resampled InferAging-BM across 30 resampling trials. The dashed line indicates the identity line (y = x). (**C**) Summary metrics for the robustness of $\delta_n^{\mathrm{BM}}$ estimates across resampling trials, including ($R^2$), MAE, and sign consistency. Each point represents one resampling trial. (**D**) Pooled comparison of biological age $z_n^{BM}$ estimates between full-data InferAging-BM and age-imbalance-resampled InferAging-BM. The dashed line indicates the identity line (y = x).

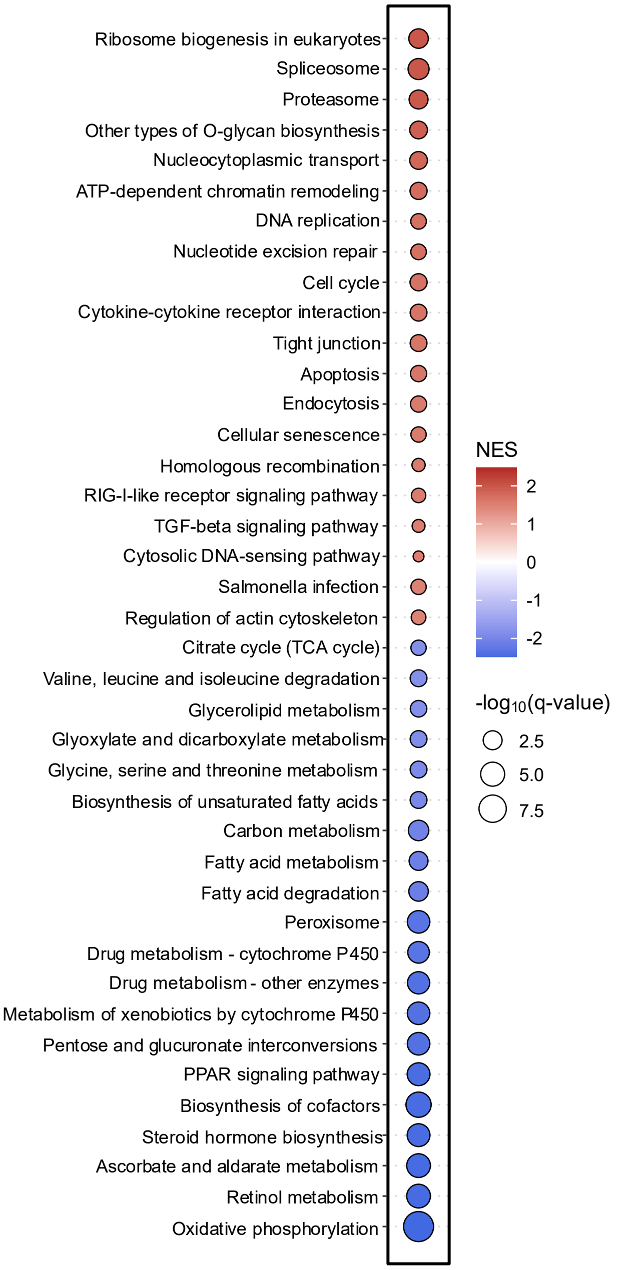


**Fig. S11. GSEA-enriched pathways associated with $z_n^{\mathrm{BM}}$.**
Dot plot of all significant pathways (q < 0.10) identified by GSEA of KEGG pathways. Genes were ranked by Spearman's rank correlation coefficients with $z_n^{\mathrm{BM}}$. The color bar indicates the normalized enrichment score (NES). The size of each dot represents $-\log_{10}$(q-value).

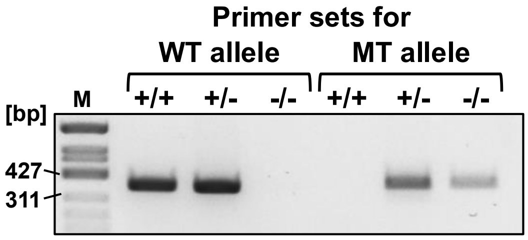


**Fig. S12. Genotyping of *kl* mutant zebrafish by allele-specific PCR.**
A 1.5% agarose gel electrophoresis image of allele-specific PCR products amplified from genomic DNA extracted from zebrafish caudal fins. Wild-type (WT) and mutant (MT) alleles were amplified using independent primer sets and cycling conditions. The genotype labels +/+, +/-, and -/- indicate wild-type, heterozygous, and homozygous mutant genotypes at the *kl* locus, respectively. Specific bands corresponding to the wild-type and mutant alleles were detected at 365 bp and 375 bp, respectively. M indicates the DNA size marker, Marker 9: $\phi$X174/HinfI digest (NIPPON GENE).

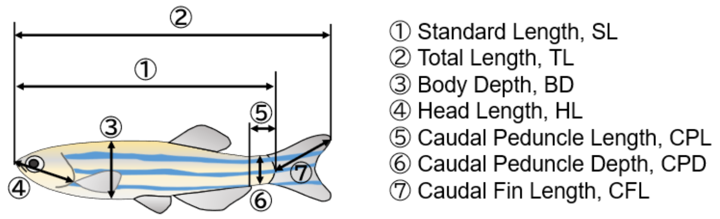


**Fig. S13. Definition of morphological landmarks in zebrafish.**
Measurement points for each morphological feature in anesthetized zebrafish. The following raw measurements were extracted using the image analysis software Fiji: SL (standard length), TL (total length), BD (body depth), HL (head length), CPL (caudal peduncle length), CPD (caudal peduncle depth), and CFL (caudal fin length).

**Supplemental table 1. Summary of statistical analyses of age-associated changes in behavioral and morphological indices**

| Feature | Kruskal–Wallis test (p-value) | Significant pairs (Dunn's test, q-value < 0.05) |
|---|---|---|
| **Variance (x)** | 0.432 | None |
| **Variance (y)** | 0.741 | None |
| **Mean of coordinates (x)** | 0.450 | None |
| **Mean of coordinates (y)** | 0.706 | None |
| **Travel distance** | 0.00401 | 4mpf vs 7mpf (q= 0.009), 4mpf vs 8mpf (q= 0.009) |
| **Travel distance (x)** | 0.00415 | 4mpf vs 7mpf (q = 0.011), 4mpf vs 8mpf (q= 0.011) |
| **Travel distance (y)** | 0.0130 | 4mpf vs 7mpf (q= 0.010) |
| **Count (Top)** | 0.879 | None |
| **Count (Middle)** | 0.228 | None |
| **Count (Bottom)** | 0.554 | None |
| **Latency to Top** | 0.921 | None |
| **Invasion to Top** | 0.0536 | 4mpf vs 7mpf (q= 0.042) |
| **Velocity (Travel distance / sec.)** | 0.00390 | 4mpf vs 7mpf (q= 0.006), 4mpf vs 8mpf (q= 0.006) |
| **Variance of Velocity** | 0.00741 | 4mpf vs 7mpf (q= 0.018), 4mpf vs 8mpf (q= 0.014) |
| **Weight** | 0.0700 | None |
| **SL_Standard Length** | 0.0879 | None |
| **TL_Total Length** | 0.0670 | None |
| **BD_Body Depth** | 0.155 | None |
| **CPL_Caudal Peduncle Length** | 0.947 | None |
| **CPD_Caudal Peduncle Depth** | 0.0317 | None |
| **HL_Head Length** | 0.0947 | None |
| **CFL_Caudal Fin Length** | 0.0749 | None |
| **BD/SL** | 0.494 | None |
| **CPD/CPL** | 0.278 | None |
| **HL/SL** | 0.939 | None |
| **Weight/TL_Total Length** | 0.0882 | None |

### Supplemental table 2. Original data for Fig. 5.

| Feature | Mean weight [95% CI] |
|---|---|
| **Variance (x)** | -0.273 [-0.703, 0.140] |
| **Variance (y)** | 0.000309 [-0.414, 0.413] |
| **Mean of coordinates (x)** | 0.0535 [-0.359, 0.473] |
| **Mean of coordinates (y)** | -0.115 [-0.532, 0.295] |
| **Travel distance** | -0.750 [-1.21, -0.333] |
| **Travel distance (x)** | -0.743 [-1.20, -0.325] |
| **Travel distance (y)** | -0.754 [-1.22, -0.337] |
| **Count (Top)** | 0.203 [-0.206, 0.626] |
| **Count (Middle)** | -0.230 [-0.652, 0.183] |
| **Count (Bottom)** | -0.0877 [-0.506, 0.330] |
| **Latency to Top** | 0.126 [-0.283, 0.537] |
| **Invasion to Top** | -0.733 [-1.19, -0.316] |
| **Velocity (Travel distance / sec.)** | -0.829 [-1.29, -0.412] |
| **Variance of Velocity** | -0.737 [-1.19, -0.319] |
| **Weight** | -0.851 [-1.26, -0.494] |
| **SL_Standard Length** | -0.831 [-1.23, -0.481] |
| **TL_Total Length** | -0.901 [-1.31, -0.546] |
| **BD_Body Depth** | -0.800 [-1.19, -0.449] |
| **CPL_Caudal Peduncle Length** | 0.191 [-0.136, 0.533] |
| **CPD_Caudal Peduncle Depth** | -0.917 [-1.32, -0.561] |
| **HL_Head Length** | -0.886 [-1.29, -0.532] |
| **CFL_Caudal Fin Length** | -0.858 [-1.26, -0.506] |
| **BD/SL** | -0.659 [-1.03, -0.315] |
| **CPD/CPL** | -0.767 [-1.16, -0.422] |
| **HL/SL** | -0.132 [-0.467, 0.192] |
| **Weight/TL_Total Length** | -0.833 [-1.23, -0.478] |
| **Cluster 1** | - 0.564 [- 0.985, - 0.180] |
| **Cluster 2** | 0.0484 [- 0.336, 0.432] |
| **Cluster 3** | 0.824 [0.430, 1.282] |
| **Cluster 4** | 0.499 [0.116, 0.918] |
| **Cluster 5** | - 0.523 [- 0.945, - 0.144] |
| **Cluster 6** | 0.737 [0.347, 1.18] |
| **Cluster 7** | 0.726 [0.335, 1.17] |
| **Cluster 8** | - 0.634 [- 1.05, - 0.249] |
| **Cluster 9** | - 0.800 [- 1.25, - 0.405] |
| **Cluster 10** | 0.480 [0.0979, 0.897] |
| **Cluster 11** | 0.00707 [- 0.376, 0.392] |
| **Cluster 12** | 0.0599 [- 0.325, 0.448] |
| **Cluster 13** | - 0.204 [- 0.596, 0.174] |
| **Cluster 14** | 0.471 [0.0944, 0.880] |
| **Cluster 15** | - 0.165 [- 0.554, 0.211] |
| **Cluster 16** | 0.481 [0.105, 0.893] |
| **Cluster 17** | 0.432 [0.056, 0.839] |
| **Cluster 18** | 0.423 [0.0477, 0.835] |
| **Cluster 19** | 0.327 [- 0.0509, 0.725] |
| **Cluster 20** | 0.229 [- 0.148, 0.621] |
